# Hypercholesterolemia Impairs Clearance of Extracellular DNA and Promotes Inflammation and Atherosclerotic Plaque Progression

**DOI:** 10.1101/2020.10.05.308478

**Authors:** Umesh Kumar Dhawan, Purbasha Bhattacharya, Sriram Narayanan, Vijayprakash Manickam, Ayush Aggarwal, Manikandan Subramanian

## Abstract

Defects in clearance of extracellular DNA due to sub-optimal activity of DNase results in exacerbated inflammation and contributes to the pathophysiology of atherosclerosis and other inflammatory diseases. However, the physiological mechanisms that regulate systemic DNase levels and the basis of its functional impairment during disease are poorly understood. Using a mouse model of experimental increase in systemic extracellular DNA levels, we identify the existence of a physiologic DNA-induced DNase response. Importantly, hypercholesterolemia in mice impairs this critical DNA-induced DNase response through an endoplasmic reticulum stress-mediated mechanism with consequences in advanced atherosclerotic plaque progression including increased extracellular DNA accumulation, exacerbated inflammation, and development of pathological features of necrotic rupture-prone vulnerable plaques. From a translational standpoint in humans, we demonstrate that individuals with hypercholesterolemia have elevated systemic extracellular DNA levels and decreased plasma DNase activity. These data suggest that the restoration of DNA-induced DNase response could be a potential therapeutic strategy to promote inflammation resolution during hypercholesterolemia.

## Introduction

Extracellular DNA released by NETosing neutrophils and necrotic cells represent a potent damage associated molecular pattern which elicits a robust inflammatory response that is critical for host defense against pathogens^1^. However, inappropriate release and persistence of extracellular DNA has been demonstrated to be maladaptive in cancers, autoimmune disorders, and several chronic inflammatory diseases including cardiometabolic disease such as atherosclerosis^2^. Therefore, prompt clearance of extracellular DNA is a critical factor to prevent host tissue damage and maintain functional homeostasis. In this context, it is important to note that the clearance of extracellular DNA is facilitated by enzymatic digestion mediated by secreted endonucleases such as DNase1 and DNase1L3^3^.

Interestingly, exogenous systemic administration of DNase1 can lower inflammation and prevent pathological progression of experimental disease settings in the mouse including breast cancer, lupus, lung injury, sepsis, and atherosclerosis^4^. These data suggest that impairment in the process of clearance of extracellular DNA can exacerbate disease progression and restoration of DNA clearance could be efficiently harnessed for therapy.

In atherosclerosis, NETotic cells and NET-DNA accumulate in atherosclerotic lesions which is causally implicated in plaque progression^5–7^. The increase in atherosclerotic plaque NET-DNA has been attributed to hypercholesterolemia-induced neutrophilia^8^ and enhanced susceptibility of neutrophils to NETosis^9^. However, whether the persistence of NET-DNA in the atherosclerotic lesions could additionally be due to impairment in DNase-mediated clearance of NET-DNA has not been explored.

Interestingly, in clinical conditions associated with extensive necrosis or NETosis such as myocardial infarction and sepsis, the levels of extracellular DNA as well as plasma DNase activity are known to increase^10–13^. Importantly, patients with myocardial infarction who had a lower ratio of ds-DNA/DNase activity had significantly better survival and recovery^14^ suggesting an important role for elevated level of DNase in attenuating disease pathogenesis. However, the mechanism of increase in DNase activity and whether it is mediated in response to an increase in the level of extracellular DNA is currently unknown.

In this study, we report that experimental increase in extracellular DNA levels leads to a concomitant increase in serum DNase activity indicating the operation of a robust DNA-induced DNase response. Most importantly, we demonstrate that hypercholesterolemia impairs this DNA-induced DNase response by an ER-stress mediated mechanism which may account for the persistent elevation of extracellular DNA levels, increased local and systemic inflammation, and advanced atherosclerosis progression. These data provide novel insights into additional mechanisms by which hypercholesterolemia impairs inflammation resolution and promotes the development of rupture-prone vulnerable atherosclerotic plaques that result in myocardial infarction and stroke.

## Results

### Serum DNase1 and DNase1L3 levels are regulated in response to changes in extracellular DNA concentration

We reasoned that the basal level of DNases in serum may be sufficient under physiological conditions but these may be regulated under conditions of inflammation that elevate circulating extracellular DNA, as contributed by dying cells such as NETosing neutrophils and necrotic cells. To directly test this hypothesis, we experimentally increased the circulating levels of extracellular DNA in mice via intravenous injection of neutrophil extracellular trap (NET) DNA which were generated from PMA-treated neutrophil-like differentiated HL-60 cells^15^ (**Sup Figure 1A and B**). First, we confirmed that intravenous injection of NET-DNA led to a rapid increase in the concentration of serum extracellular ds-DNA (**Figure 1A**). Interestingly, the concentration of extracellular DNA decreased over time and returned to homeostatic levels within 12 h (**Figure 1A)**indicating the operation of an efficient extracellular DNA clearance mechanism. Next, we asked whether the return to homeostatic levels of extracellular DNA involved an increase in the level of serum DNases. To address this question, we conducted an *in-vitro* DNA degradation assay by incubating a defined quantity of genomic DNA with serum isolated from mice injected vehicle or NET-DNA. Interestingly, serum from NET-DNA injected mice degraded ~80% of the input DNA as compared with only ~20% degradation observed in vehicle-injected mice (**Figure 1B**). These data suggested that total DNase activity is higher in the serum of DNA-injected mice. To quantify the increase in DNase activity, we conducted Single Radial Enzyme Diffusion (SRED) assay as described previously^3^ where the area of DNA degradation is a measure of the absolute units of DNase activity as measured using standards of known concentration of DNase. Consistent with the *in-vitro* DNA degradation assay, we observed that NET-DNA-injected mice demonstrated a ~3-fold increase in serum total DNase activity at 12 h post-injection of DNA (**Figure 1C**). Next, to understand the dynamics of the increase in DNase activity in response to extracellular DNA, we conducted a SRED assay and quantified the serum total DNase activity at 1-, 4-, 8-, and 12-h post-injection of NET-DNA. A significant increase in serum total DNase activity was quantified as early as 4 h post-injection of DNA and reaching a peak at 8 h indicating a rapid active regulation of serum DNase in response to an increase in extracellular DNA (**Figure 1D**).

**Figure 1.**
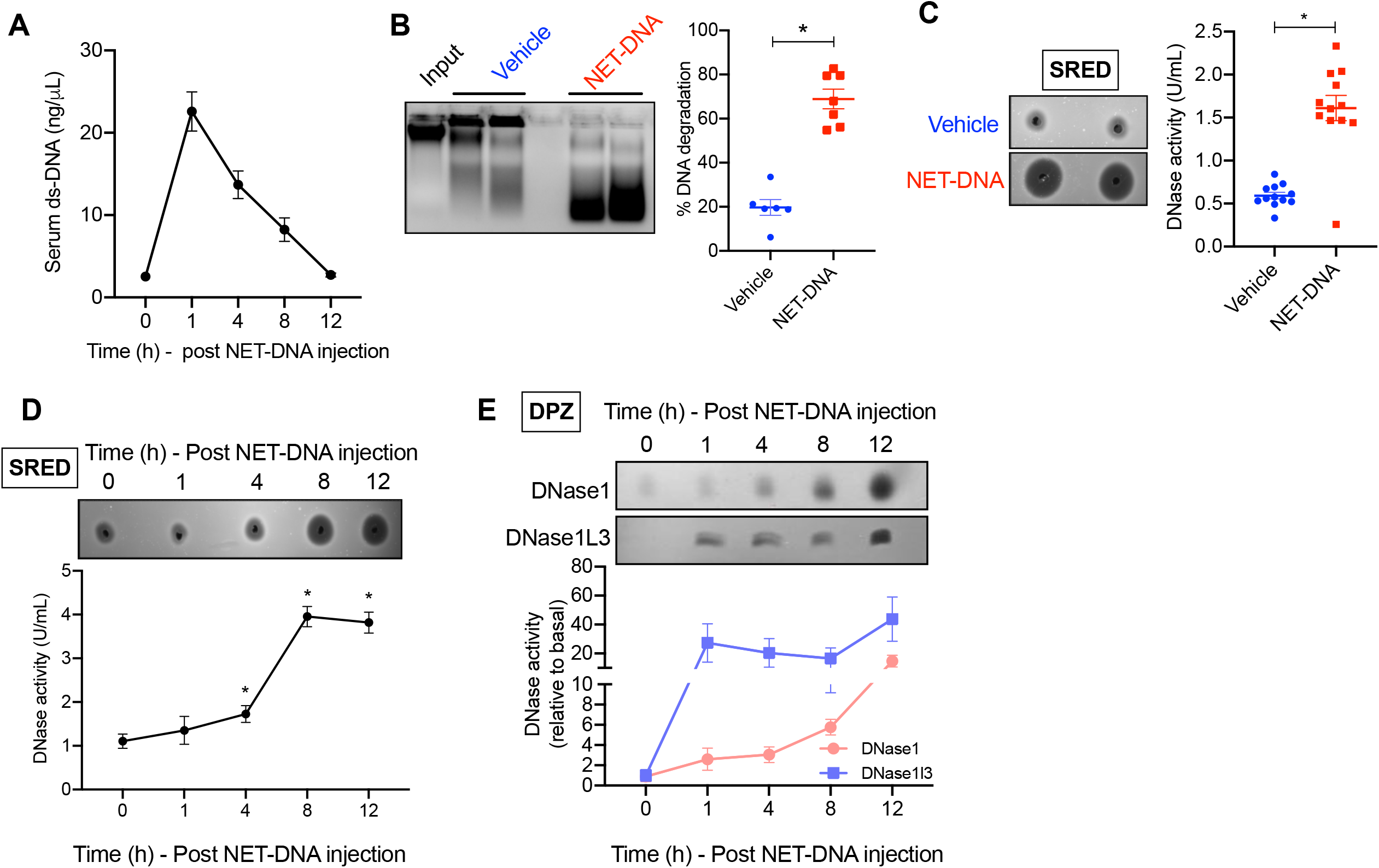
Increase in extracellular DNA levels elicits a systemic DNase response. WT mice were administered NET-DNA (40 g) intravenously followed by collection of blood at periodic intervals as indicated. **(A)** Sera of mice were analyzed for the levels of extracellular ds-DNA using Qubit fluorometric assay. **(B)** Sera collected at 12 h from vehicle or NET-DNA-injected mice were incubated with input DNA for 2.5 h followed by agarose gel electrophoresis. The scatter plot on the right represents quantified data of the extent of DNA degradation expressed as a percentage of the input DNA (n= 5 mice in vehicle-treated and 6 mice in NET-DNA treated group). **(C)** Single radial enzyme diffusion (SRED) assay to quantify total DNase activity in the serum collected at 12 h from vehicle or NET-DNA-injected mice. The scatter plot represents the quantified data of DNase activity in the two groups of mice obtained by interpolation from known DNase standards. n = 12 mice in each group. **(D)** Similar to (C) except that SRED was conducted on sera collected from NET-DNA injected mice at indicated time points. **(E)** Depolymerizing polyacrylamide gel electrophoresis zymography (DPZ) assay to analyze expression levels of DNase1 and DNase1L3 in the sera of NET-DNA injected mice at the indicated time points. n = 4 mice per group. *, p < 0.05 by Mann-Whitney test.

An increase in serum DNase activity could be mediated by either an increase in the level of DNases in serum or an increase in activity without actual changes in the expression level due to a decrease in the levels of endogenous plasma DNase inhibitors such as heparin, plasmin, and aprotinin^16^. To understand whether the increase in serum DNase activity is mediated by an actual increase in DNase levels, we conducted Depolymerizing Gel Zymography (DPZ) as described previously^3,16^. In this assay, DNase in the serum is denatured and resolved on a polyacrylamide gel followed by in-gel protein refolding to recover DNase activity. This assay thereby removes the confounding effects of DNase inhibitors present in serum and measures the true serum DNase levels as a function of DNase activity. Furthermore, by using specific denaturing and renaturing conditions during DPZ, the levels of the two major serum DNases, namely, DNase1 and DNase1L3 can be independently quantified^3,16^. DPZ demonstrated that the increase in DNase activity upon NET-DNA injection was contributed by an increase in the levels of both DNase1 and DNase1L3 (**Figure 1E**). In summary, these data demonstrate that the level of key serum DNases, DNase1 and DNase1L3, are regulated in response to an increase in extracellular DNA presumably to degrade and clear extracellular DNA and restore homeostasis.

### Hypercholesterolemia leads to elevated extracellular DNA and impaired DNase response

The data presented above suggest that extracellular DNA levels are tightly regulated by changes in DNase expression to enable maintenance of homeostasis. In this context, it is interesting to note that persistence of NET- and necrotic cell DNA has been reported in atherosclerotic plaques^6–8^ which raises the interesting possibility that the clearance of extracellular DNA may be impaired in atherosclerosis as well as other pathological conditions.

Since hypercholesterolemia is one of the initiator and driver of atherosclerosis^17^, we tested whether hypercholesterolemia could impair systemic extracellular DNA clearance. Towards this end, we set up an animal model of normocholesterolemia (WT *C57BL6* mice on chow diet), moderate hypercholesterolemia (*Apoe*^*−/−*^ mice on chow diet), and severe hypercholesterolemia (*Apoe*^*−/−*^ mice on a western type diet for 3 weeks) (**Figure 2A**). Interestingly, we observed that under basal conditions, the level of serum extracellular DNA was significantly higher in moderate and severely hypercholesterolemic mice as compared with normocholesterolemic mice (**Figure 2B**). In addition, there was a significant positive correlation between plasma total cholesterol levels and plasma extracellular DNA levels in this experimental model of hypercholesterolemia (**Figure 2C**). Since, the levels of extracellular DNA is regulated by DNase activity, we next asked if a decrease in the level of plasma DNase could account for the increase in ds-DNA observed during hypercholesterolemia. Surprisingly, we observed that the basal DNase activity was higher in mice with moderate hypercholesterolemia as compared with normocholesterolemic mice while the total DNase activity of severely hypercholesterolemic mice was comparable with control mice (**Figure 2D**). These data in conjunction with the observation of persistently high plasma extracellular DNA levels in hypercholesterolemic mice raised the possibility that the DNA-induced DNase response might be sub-optimal during hypercholesterolemia. To test this hypothesis and to quantify the DNA-induced DNase response, we injected NET-DNA systemically in these mice and measured the rate of clearance of extracellular DNA from the circulation as well as the upregulation of DNase activity in the serum. We observed that the moderately hypercholesterolemic mice demonstrated a DNA-induced DNase response which was comparable to control mice **(Figure 2E)**. However, the severely hypercholesterolemic mice demonstrated a significantly blunted DNA-induced DNase response upto the 12 h period of observation and was lower by about 50% as compared with the other groups of mice (**Figure 2E)**. Next, we conducted DPZ which demonstrated that the decrease in total DNase activity in the severely hypercholesterolemic mice was due to a decrease in the release of both DNase1 and DNase1L3 (**Figure 2F)**. Since there is a decrease in the total DNase activity in the severely hypercholesterolemic mice, we asked whether this results in impairment in the rate of clearance of injected NET-DNA. Compared with control and moderately hypercholesterolemic mice, the severely hypercholesterolemic mice had significant delay in the clearance of NET-DNA (**Figure 2G)**. Next, we measured the resolution interval^18^ which we have defined as the time taken for extracellular DNA level to reach half-maximum as an indicator of efficiency of extracellular DNA clearance and return to homeostasis. The resolution interval in the severely hypercholesterolemic mice was almost double the control mice suggesting impaired clearance of extracellular DNA and delay in restoration of homeostasis (**Figure 2H)**.

**Figure 2.**
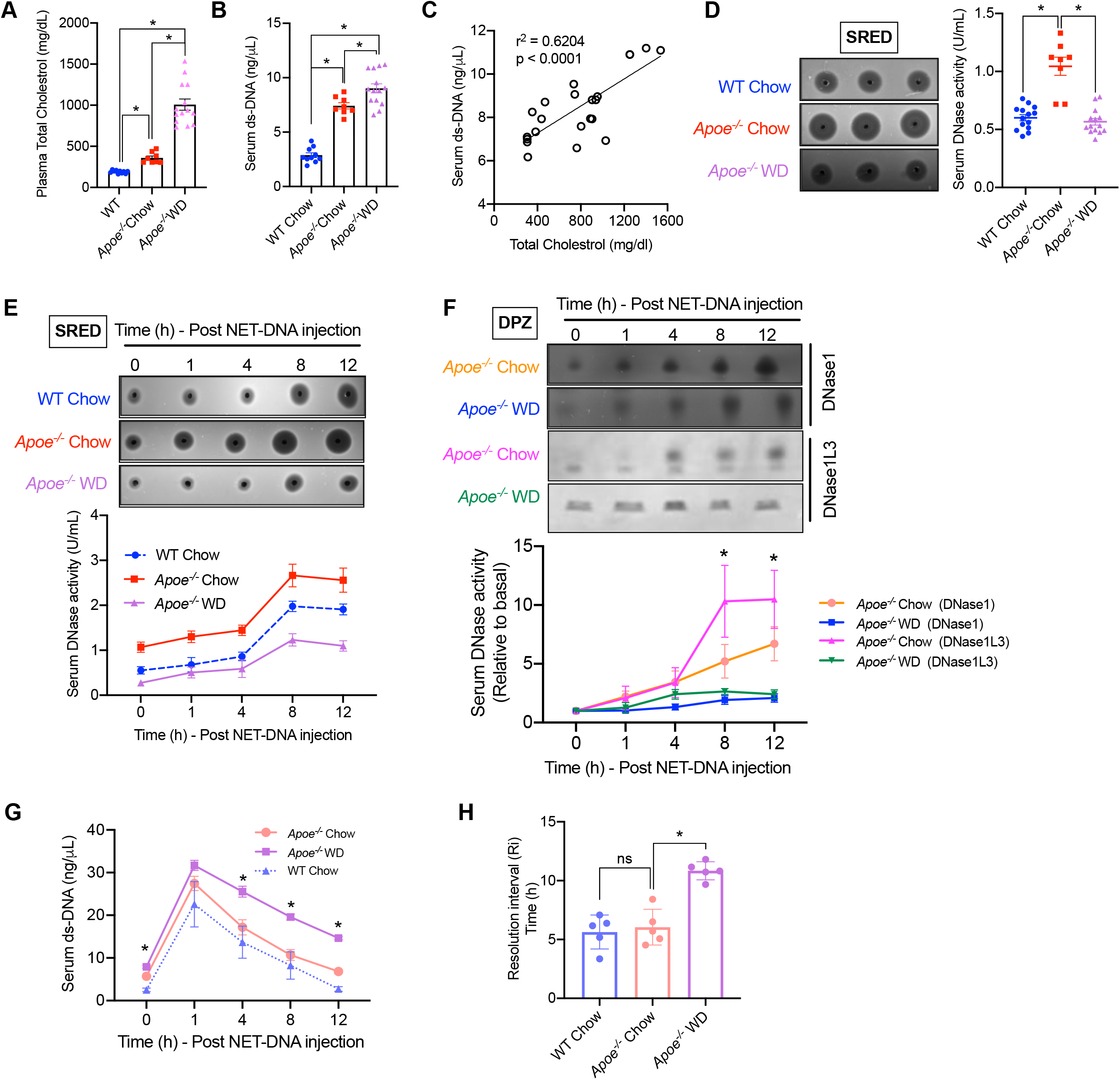
Hypercholesterolemia impairs the DNA-induced DNase response. 8 wk-old WT C57BL/6 mice and *Apoe*^*−/−*^ mice were fed either a standard chow diet (chow) or a western-type diet (WD) for 3 wks as indicated. **(A)** Measurement of plasma total cholesterol in indicated groups of mice. n = 8-14 mice per group. **(B)** The serum extracellular ds-DNA levels were measured in the indicated groups of mice using the Qubit fluorescence assay (n = 8-14 mice group). **(C)** The plot represents correlation between total cholesterol levels and serum extracellular ds-DNA concentration. Linear regression was conducted to analyze correlation between the two parameters. **(D)** Quantification of basal total serum DNase activity by SRED in WT and *Apoe*^*−/−*^ mice after 3 wks on indicated diet. **(E-H)** The three groups of mice were administered NET-DNA intravenously and sera was isolated at periodic intervals as indicated and subjected to analysis of **(E)** total serum DNase activity by SRED, **(F)** analysis of DNase1 and DNase1L3 activity by DPZ, **(G)** analysis of extracellular DNA levels, and **(H)** analysis of resolution interval defined as the time taken for the extracellular DNA levels to reach half-maximal. n = 5 mice per group. *, p < 0.05.

Next, we addressed whether the hypercholesterolemia-induced decrease in DNase response and increase in extracellular DNA are observed in humans. Interestingly, we observed that individuals with hypercholesterolemia (plasma total cholesterol > 200 mg/dL) had significantly higher levels of plasma extracellular DNA as compared with normocholesterolemic individuals (**Figure 3A)**. In addition, there was a significant positive correlation between the levels of plasma total cholesterol and plasma extracellular DNA (**Figure 3B)** similar to our observation in the mouse model of hypercholesterolemia. Most importantly, hypercholesterolemic individuals had significantly lower plasma total DNase activity as compared with normocholesterolemic individuals (**Figure 3C)**. Also, we observed a significant negative correlation between total plasma cholesterol and plasma total DNase activity (**Figure 3D)**. These data demonstrate that hypercholesterolemia-induced impairment of DNase activity is conserved in humans and could have pathophysiological consequences.

**Figure 3.**
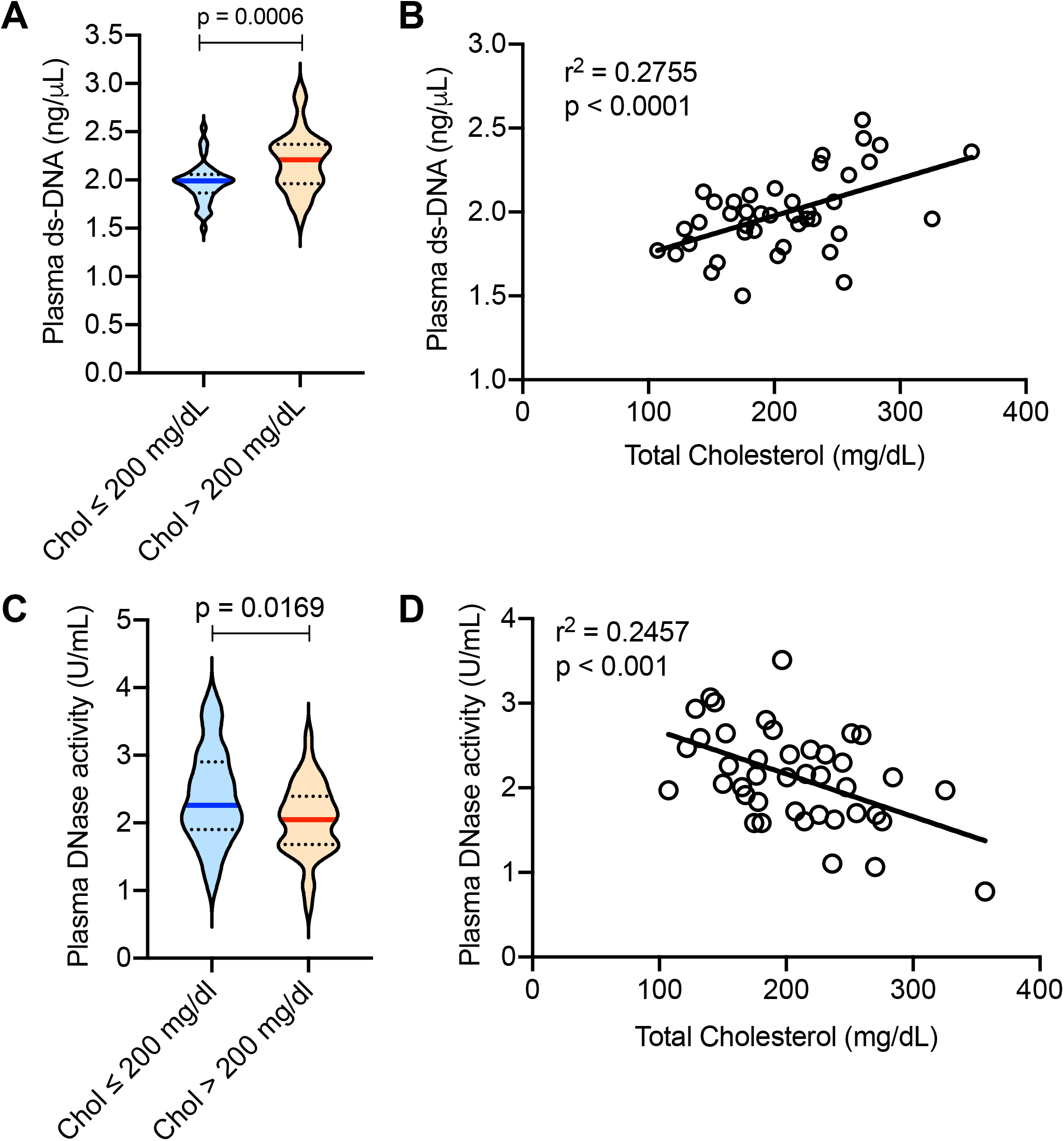
Hypercholesterolemia is associated with increased extracellular DNA and decreased plasma DNase activity in humans. Plasma was isolated from healthy adults. **(A and C)** Analysis of plasma extracellular ds-DNA levels by Qubit fluorometric assay and total DNase activity by SRED respectively in individuals with total plasma cholesterol levels 200 mg/dL (normocholesterolemia) or > 200 mg/dL(hypercholesterolemia). n = 41 per group. *, p < 0.05. **(B and D)** Analysis of correlation between total plasma cholesterol concentration with extracellular ds-DNA levels or total DNase activity respectively. n = 41. Simple linear regression was conducted to analyze correlation.

### DNase1 treatment decreases atherosclerotic plaque extracellular DNA content, inflammation, and necrotic core formation

Previous studies have shown that accumulation of extracellular DNA, particularly NETotic, necrotic, and necroptotic DNA is involved in atherosclerotic plaque progression^6–8^. Most importantly, treatment with DNase1 has been demonstrated to prevent plaque progression in early atherosclerosis^9,19^ and promote plaque regression in diabetic mice^20^. Based on these previous studies as well as our data demonstrating impaired DNase response during hypercholesterolemia (**Figure 2E**), we next addressed the specific question whether restoration of DNase activity via exogenous supplementation of DNase1 could decrease plaque inflammation and prevent plaque progression specifically in the context of advanced atherosclerosis which has not been explored so far. To this end, we fed *Apoe*^*−/−*^ mice a high fat-high cholesterol western-type diet (WD) for 16 weeks which leads to generation of advanced atherosclerotic plaques characterized by large necrotic areas. Post 16 weeks of WD feeding, the mice were randomized into two groups with one group of mice receiving DNase1 (400 U, intravenously)^19^ three times a week for 4 weeks, while the other group was administered an equal volume of 1X-PBS as control. At the end of 4 weeks of treatment, both groups of mice had similar body weight and metabolic parameters including blood glucose, total cholesterol, and triglycerides (**Supplementary Figure 2A-D**). As expected, the plasma total DNase activity was significantly higher in mice receiving DNase1 (**Figure 4A**). Consistent with an increase in plasma DNase activity, there was a significant decrease in the plasma extracellular DNA levels in mice administered DNase1 (**Figure 4B**) reaching levels similar to that observed in *Apoe*^*−/−*^ mice on a chow diet. Most importantly, aortic root atherosclerotic lesional NET-DNA content, measured as the extent of lesions that stain positive with anti-citrullinated H3 antibody, was significantly lower in the DNase1-treated mice (**Figure 4C**). In addition, the DNase1-treated mice demonstrated a significant decrease in the plaque necrotic area (**Figure 4D**), an increase in lesional collagen content (**Figure 4E**) and smooth muscle cell numbers (**Figure 4F**), and a decrease in plaque macrophage content (**Figure 4F**). Since, the total atherosclerotic lesion area is not significantly different between the two groups of mice (**Figure 4D**), these data taken together suggest that DNase1 treatment leads to significant cellular and structural remodeling of atherosclerotic lesions towards a stable-plaque phenotype. Importantly, DNase1 treatment led to a significant decrease in atherosclerotic plaque inflammation as demonstrated by the decreased expression of pro-inflammatory genes such as *Tnf*, *Ifng*, *Ifnb*, and *Mx1* (**Figure 4G**). Also, DNase1-treated mice had lower systemic inflammation as demonstrated by decreased levels of pro-inflammatory gene expression in spleen (**Supplementary Figure 2E**) and lower levels of several pro-inflammatory cytokines in the plasma (**Figure 4H**). These data suggest that therapeutic increase in DNase activity to overcome the sub-optimal DNA-induced DNase response in hypercholesterolemia leads to beneficial effects in promoting plaque stability and lowering systemic inflammation.

**Figure 4.**
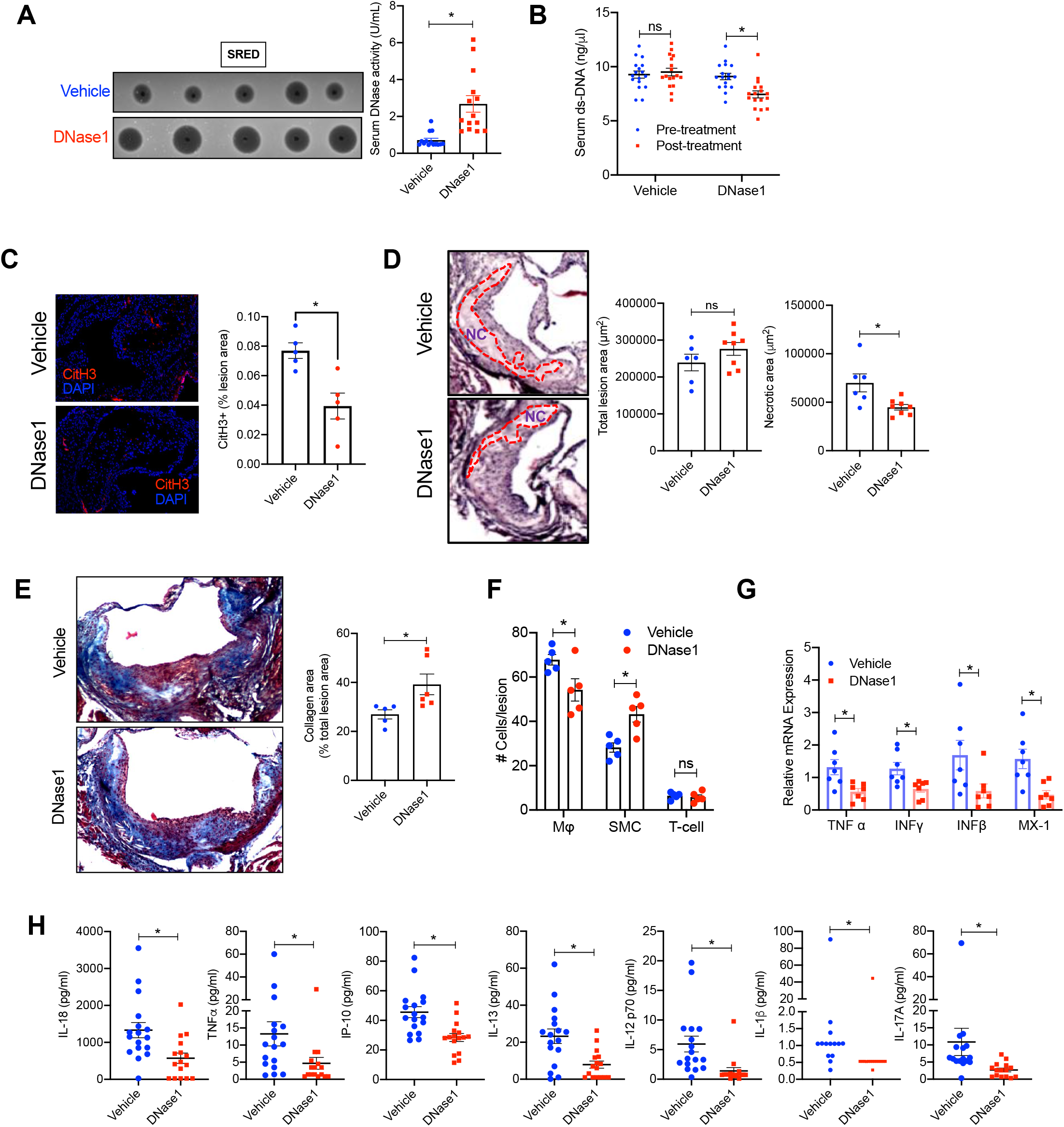
Exogenous DNase1 administration promotes atherosclerotic plaque remodeling and stabilization. *Apoe^−/−^* mice were fed western-type diet for 16 wks followed by administration of either vehicle or DNase1 (400 U) intravenously 3 times a week for 4 weeks. **(A)** SRED analysis of serum total DNase activity in vehicle or Dnase1-injected mice. **(B)** Analysis of serum extracellular ds-DNA concentration in indicated groups of mice before and after treatment with vehicle or DNase1. **(C)** Aortic root sections of vehicle and DNase1-treated mice were immunostained with antibody against CitH3 (red) and percent lesion area that stains positive for CitH3 was quantified. Nucleus were stained with DAPI (blue). n = 5 mice per group. **(D)** H&E staining of aortic root sections of vehicle and DNase1-treated mice to quantify the total lesion area and the total necrotic area (red dotted line). **(E)** Mason trichrome staining of aortic root sections of vehicle and DNase1-treated mice to quantify collagen deposition in the plaque. **(F)** Immunofluorescence-based quantification of percent macrophages (F4/80), smooth muscle cells (sm-Actin), and T-cells (CD3) in atherosclerotic lesions of vehicle and DNase1-treated mice. **(G)** qPCR-based analysis of relative mRNA expression of indicated pro-inflammatory genes in aorta of vehicle and DNase1-treated mice. **(H)** Multiplex ELISA for quantification of pro-inflammatory cytokine levels in the sera of vehicle and DNase1-treated mice. The data are represented as mean SEM. *, p < 0.05; ns, no significant difference. Mann-Whitney test was conducted to analyze statistical significance.

### Macrophage-mediated clearance of extracellular DNA is mediated by secretion of DNase1 and DNase1L3

In theory, it is possible that the impaired clearance of extracellular DNA in the atherosclerotic plaque is mediated by a decrease in lesional DNase activity secondary to the sub-optimal levels of DNase in circulation during hypercholesterolemia. However, it is known that myeloid cells, particularly macrophages and dendritic cells, are one of the major source of secreted DNase1L3^21^ raising the possibility that the impaired clearance of extracellular DNA in the atherosclerotic plaque could potentially be mediated by a defect in the local secretion of DNases by lesional macrophages. Indeed, our analysis of publicly available RNA-sequencing data of atherosclerotic plaque macrophages^22^ revealed that they express transcripts for both DNase1 and DNase1L3 (**Supplementary Figure 3A**). Most importantly, immunofluorescence analysis of atherosclerotic lesions from 16 wk WD-fed *Apoe*^*−/−*^ mice demonstrated that plaque macrophages stain positive for intracellular DNase1 and DNase1L3 (**Figure 5A**) suggesting that plaque macrophages could be a local source of DNases.

**Figure 5.**
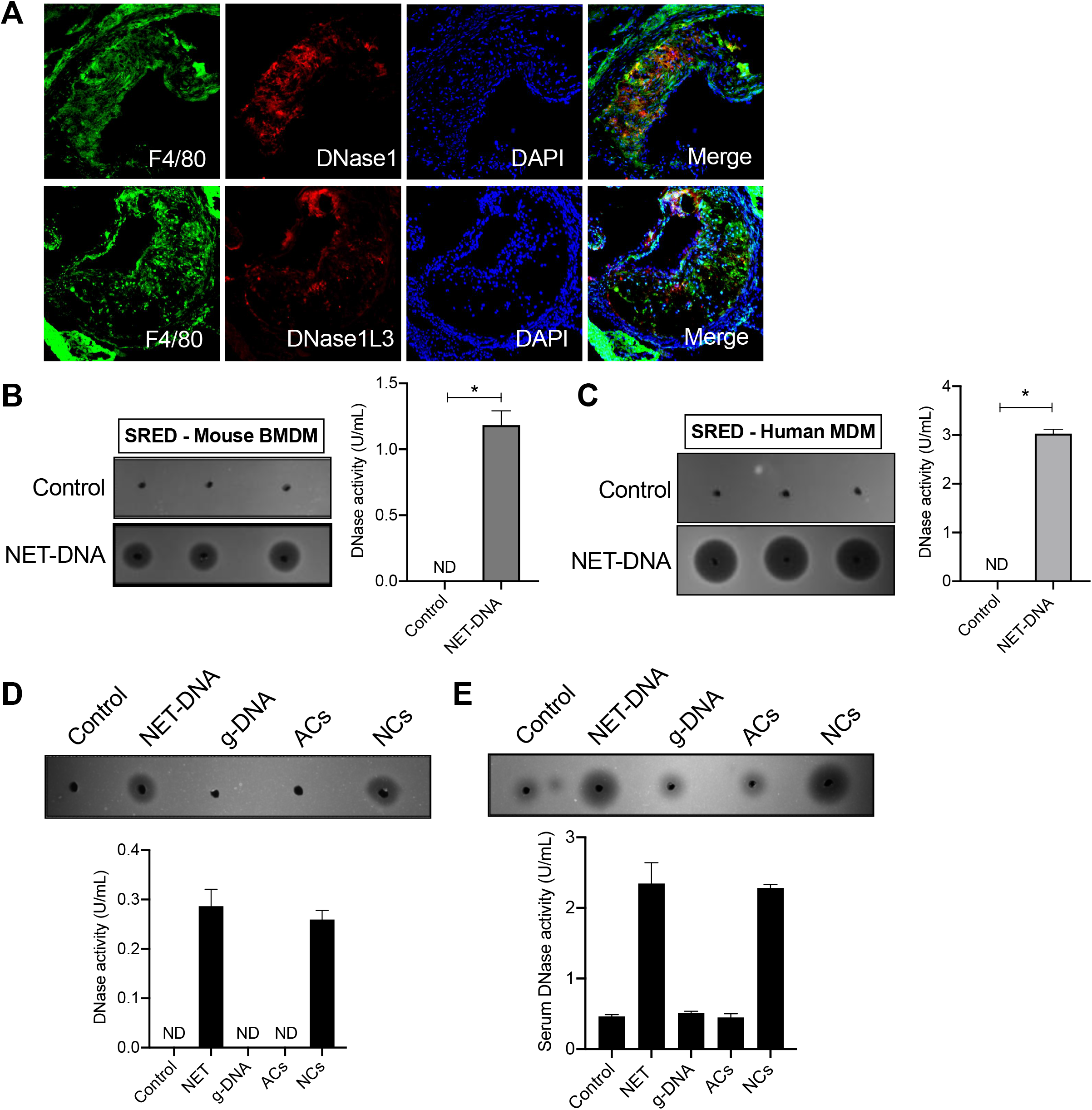
Macrophages secrete DNases in response to extracellular DNA. **(A)** Aortic root sections of 16 wk WD-fed *Apoe*^*−/−*^ mice were immunostained with anti-DNase1 antibody (top panel) or anti-DNase1L3 antibody (bottom panel) and their expression colocalized with macrophages stained with anti-F4/80 antibody (green). Nucleus was counterstained with DAPI (blue). **(B)** SRED assay for quantification of total DNase activity in the culture supernatants of murine BMDM either left untreated or treated with NET-DNA (10 g) for 12 h. **(C)** SRED assay for quantification of total DNase activity in the culture supernatants of human peripheral blood monocyte-derived macrophages either left untreated or treated with NET-DNA for 12 h. **(D)** SRED assay for quantification of total DNase activity in culture supernatants of murine BMDMs treated for 12 h with either NET-DNA (10 g), genomic DNA (g-DNA, 10 g), apoptotic cells (ACs, 1:5 macrophage:AC ratio), or necrotic cells (NCs, 1:5 macrophage:NC ratio) as indicated. **(E)** SRED-based determination of total DNase activity in the serum of mice injected with NET-DNA, g-DNA, ACs, or NCs.

Although, macrophages express transcripts for DNase1 and DNase1L3 and show intracellular protein expression, consistent with previous literature^23^, we observed that *in-vitro* macrophages under basal conditions secrete very low quantities of DNase1 and DNase1L3 (**Supplementary Figure 3B**). It is important to note that this basal quantity of secreted DNase was detected by DPZ after concentration of macrophage culture supernatant. These levels were insufficient to detect any measurable DNase activity in either SRED or the *in-vitro* DNA degradation assay (**Supplementary Figure 3C-D**). In this context, we asked whether macrophages exposed to NET-DNA have the ability to sense them and respond via enhanced secretion of DNase1 or DNase1L3. Towards this end, we incubated murine bone marrow-derived macrophages (BMDMs) with NET-DNA for 12 h and quantified DNase activity in the supernatant by SRED. As shown in **Figure 5B**, supernatant from BMDMs incubated with NET-DNA demonstrated a significant increase in DNase activity indicating macrophage-mediated secretion of DNases. Indeed, a similar response was observed in human THP-1 macrophage-like cells (**Supplementary Figure 3E**) and human peripheral blood monocyte-derived macrophages (**Figure 5C**) which clearly establishes that both murine and human macrophages have the ability to sense extracellular DNA and respond via secretion of DNases.

Since NET-DNA is composed of DNA in complex with several proteins including components of the chromatin, we next asked whether the DNase response is mediated by sensing DNA or some component of the DNA-protein complex. Interestingly, DNase was released only when macrophages were incubated with NET-DNA but not with genomic DNA that is stripped of its associated proteins (**Figure 5D**). In addition, DNase release was also observed when macrophages were incubated with necrotic cells which are known to release nuclear DNA upon loss of membrane integrity (**Figure 5D**). In contrast to necrotic cells, incubation of macrophages with apoptotic cells did not elicit a DNase response (**Figure 5D**). Importantly, similar data were obtained *in-vivo* wherein mice injected either NET-DNA or necrotic cells demonstrated a robust increase in plasma total DNase activity, while injection of genomic DNA or apoptotic cells did not elicit a DNase response (**Figure 5E**). In summary, these data suggest that the DNase response is specific and mediated by cellular sensing of a DNA-protein complex.

Next, we asked the question whether the increase in DNase activity in culture supernatants of macrophages incubated with NET-DNA is mediated via secretion of DNase1 or DNase1L3. DPZ of culture supernatant from BMDMs incubated with NET-DNA revealed that macrophages secrete both DNase1 and DNase1L3 (**Figure 6A**). Similar to our *in-vivo* observation where the levels of plasma total DNase increases significantly within 4 h of NET-DNA injection (**Figure 1D**), macrophages exposed to NET-DNA led to a very rapid increase in the levels of DNase1 and DNase1L3 in the culture supernatant (**Figure 6B**). Interestingly, we observed that BMDMs incubated with NET-DNA did not show changes in the expression levels of DNase1 or DNase1L3 mRNA (**Supplementary Figure 4A**) indicating a lack of transcriptional regulation in this process. Consistent with this observation, our analysis of publicly available data^20^ revealed no significant differences in the mRNA expression levels of DNase1 and DNase1L3 in plaque macrophages that are in the proximity of NET-DNA+ area vs. those that were in the NET-DNA negative area (**Supplementary Figure 4B**). Additionally, macrophages pre-incubated with cycloheximide, a protein translational inhibitor^24^, were still able to increase the DNase activity in the culture supernatant upon exposure to NET-DNA (**Supplementary Figure 4C**) suggesting no requirement for protein translation in the DNase secretion process.

**Figure 6.**
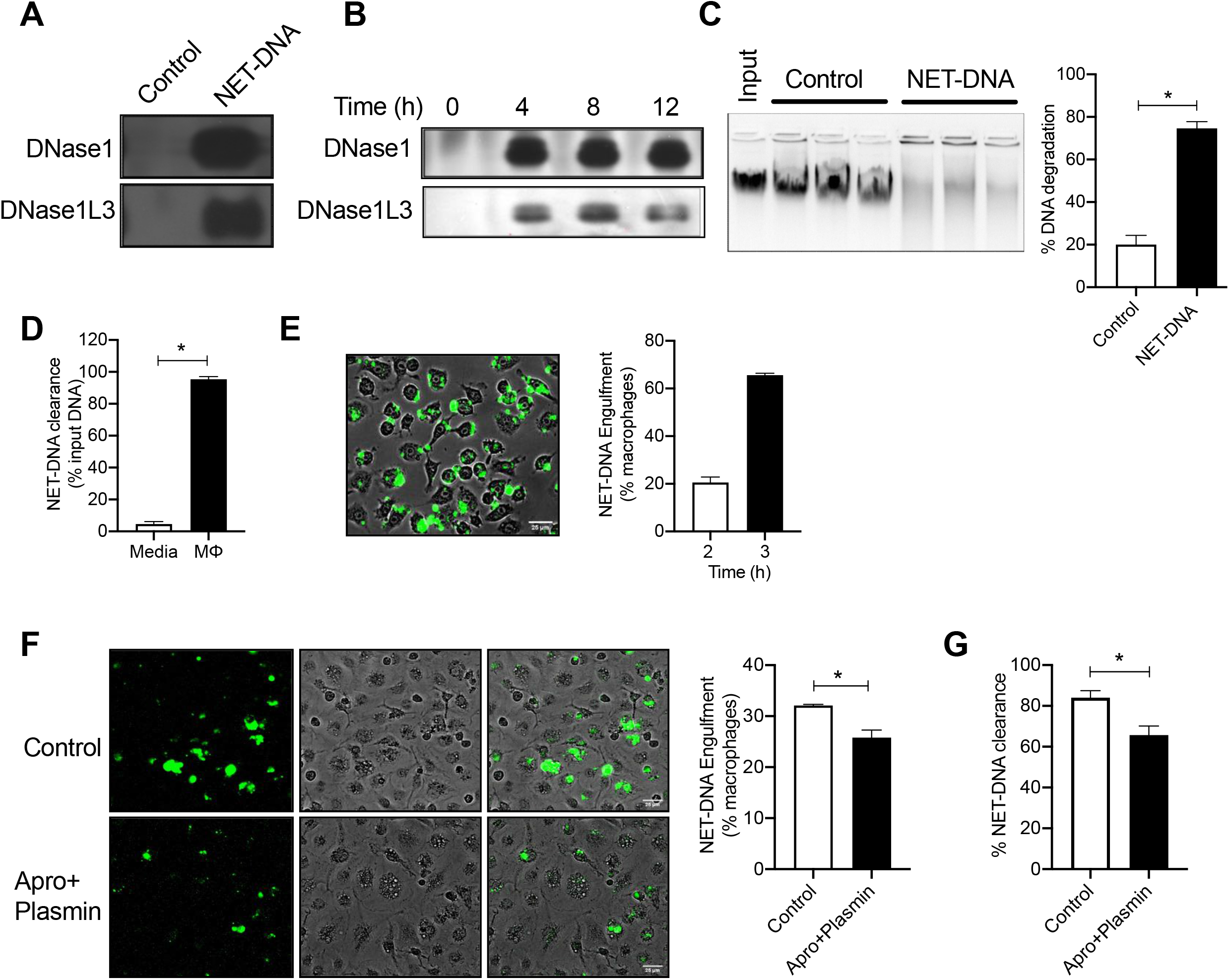
Macrophage secreted DNase1 and DNase1L3 facilitates NET-DNA engulfment and clearance. **(A)** DPZ for detection of DNase1 and DNase1L3 activity in the culture supernatants of murine BMDMs either left untreated (control) or incubated with 10 g NET-DNA for 12 h or **(B)** at indicated time points post-incubation with NET-DNA. **(C)** In-vitro DNA degradation assay as described in methods section with cell culture supernatants from BMDMs treated without or with NET-DNA. **(D)** Qubit fluorometric analysis of remaining input DNA after 12 h incubation with either cell culture media or macrophages. The data are represented as percent of input DNA that has been cleared from the media or supernatant. **(E)** Representative image demonstrating engulfment of sytox green-labeled NET-DNA (green) by BMDMs at 3 h post-incubation. The bar graph represents the percent macrophages that have engulfed NET-DNA at 2- and 3 h post-incubation. **(F)** Fluorescence microscopy-based quantification of NET-DNA engulfment efficiency of macrophages treated with vehicle or aprotinin and plasmin. **(G)** Similar to (F) except that total NET-DNA clearance at 12 h post-incubation was quantified using Qubit assay.

While it is clear from the above data that macrophages secrete DNases in response to extracellular DNA, we next asked whether the secreted DNase is relevant for macrophage mediated clearance of extracellular DNA. To address this question, we conducted an *in-vitro* DNA degradation assay by incubating a defined quantity of DNA with culture supernatant from either control macrophages or NET-DNA-exposed macrophages. We observed that culture supernatant from NET-DNA-exposed macrophages were able to efficiently degrade DNA indicating the presence of significant DNase activity (**Figure 6C**). It is important to note that the above and subsequent experiments were conducted by culturing macrophages in serum-free media for the entire duration of the experiment to remove the potential confounding issue of serum-contained DNases. Next, we incubated macrophages with NET-DNA for 12 h and measured the amount of remaining NET-DNA in the supernatant as an indicator of the efficiency of total NET-DNA clearance which is a combination of DNase-mediated extracellular degradation and macrophage-mediated engulfment of NETs followed by their intracellular degradation. Approximately, 90% of the input DNA (5 μg) was cleared from the supernatant at 12 h post-incubation with macrophages (**Figure 6D**). In addition to DNase-mediated DNA degradation, we observed that ~20% of macrophages had engulfed NET-DNA within 2 h which increased to ~ 60% at 3 h (**Figure 6E**). Interestingly, two previous studies^23,25^ have raised the possibility that DNases could aid the engulfment of DNA by macrophages via cleaving the nucleic acid into smaller fragments. However, direct physiological relevance of secreted DNase in macrophage engulfment was not adequately addressed in these studies. Hence, we asked the question, whether macrophage-secreted DNase could improve the efficiency of engulfment of NET-DNA by macrophages. Towards this end, we incubated macrophages with NET-DNA in the presence of aprotinin and plasmin, inhibitors of DNase1 and DNase1L3 respectively^16^, to inhibit the activity of macrophage-secreted DNases. Interestingly, we observed that aprotinin and plasmin treatment led to a decrease in both the percent macrophages that engulfed NET-DNA (**Figure 6F**) as well as total NET-DNA clearance (**Figure 6G**). Complementary to these findings, we observed that pre-digested NET-DNA was more efficiently engulfed by macrophages as compared with intact NET-DNA (**Supplementary Figure 4D**). These data together suggest that local secretion of DNases by macrophages promote extracellular degradation of DNA into smaller fragments which in turn enhances the ability of macrophages to efficiently engulf DNA.

### Atherogenic lipids induce ER stress and impair secretion of DNase1 and DNase1L3

Since, our *in-vivo* data demonstrates that hypercholesterolemia impairs the DNA-induced DNase response, we next asked whether macrophages exposed to hypercholesterolemic conditions have impairment in the release of DNase upon exposure to extracellular DNA. To model atherogenic dyslipidemia, we treated BMDMs with 7-ketocholesterol (7-KC), a modified oxysterol which is present abundantly in atherosclerotic plaques^26^. 7-KC treatment leads to formation of foamy macrophages similar to that observed in atherosclerotic plaques^27^. Interestingly, these 7-KC treated macrophages secreted significantly lower levels of DNase upon exposure to NET-DNA as compared with control macrophages exposed to NET-DNA (**Figure 7A**). This decrease in DNase activity was due to an impairment in the secretion of both DNase1 and DNase1L3 (**Figure 7B**). Consistent with a decrease in DNase secretion, 7-KC treated macrophages showed significantly decreased clearance of extracellular DNA (**Figure 7C**) and lower level of engulfment of NET-DNA (**Figure 7D and Supplementary Figure 5A**). It is important to note that the concentration of 7-KC used in these assays did not enhance macrophage cell death (**Supplementary Figure 5B**). Hence, the impaired DNase secretion and decreased engulfment of DNA observed in 7-KC-treated macrophages is unlikely to be due to compromised cell viability.

**Figure 7.**
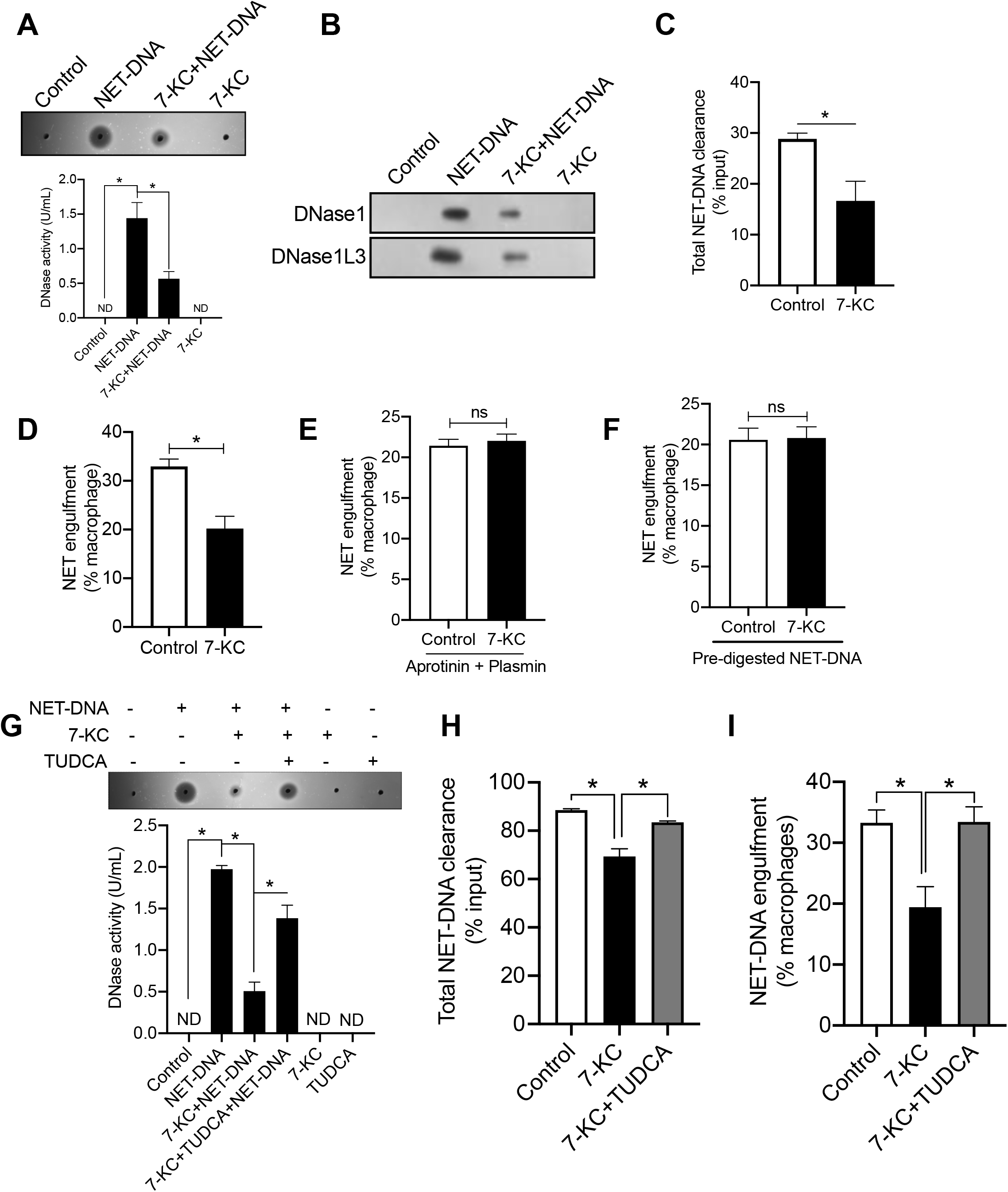
Cholesterol impairs DNA-induced DNase response in macrophages via ER stress. **(A)** SRED for quantification of total DNase activity and **(B)** zymography for detection of DNase1 and DNase1L3 activity in the cell culture supernatants of the indicated group of BMDMs. **(C)** Qubit fluorometric analysis of the total NET-DNA clearance efficiency and **(D)** NET-DNA engulfment efficiency of control and 15 M 7-KC-treated macrophages. **(E)** Fluorescence microscopy-based analysis of the efficiency NET-DNA engulfment by control and 7-KC-treated macrophages in the presence of Dnase inhibitors aprotinin (100 U/mL) and plasmin (40 U/mL). **(F)** Analysis of the engulfment efficiency of control and 7-KC-treated macrophages incubated with pre-digested NET-DNA. **(G)** SRED-based analysis of total DNase activity in cell culture supernatants of indicated groups of macrophages. **(H)** Qubit fluorometric analysis of the total NET-DNA clearance efficiency and **(I)** NET-DNA engulfment efficiency of indicated group of macrophages. The data are representative of three independent experiments. Student’s t-test or ANOVA was conducted on appropriate groups to determine statistical significance. *, p < 0.05. ns, no statistical significance.

Since, as shown above, the efficiency of engulfment of extracellular DNA is influenced by DNase-mediated degradation, it is possible that the reduced engulfment efficiency of 7-KC-treated macrophages is due to a decrease in secretion of DNases. Hence, we set-up an experiment to address the question whether 7-KC-treated macrophages have an additional defect in engulfment of extracellular DNA that is independent of decreased DNase-mediated degradation of DNA. Interestingly, we observed that the efficiency of engulfment of extracellular DNA of 7-KC-treated macrophages was similar to that of control macrophages when the activity of DNase1 and DNase1L3 were inhibited with aprotinin and plasmin respectively (**Figure 7E**). In addition, the extracellular DNA engulfment efficiency of 7-KC-treated macrophages was similar to that of control macrophages when they were incubated with pre-digested DNA. (**Figure 7F**). These data together suggest that the defect in extracellular DNA engulfment observed in 7-KC-treated macrophages was primarily due to the decrease in secretion of DNases which impairs its ability to fragment NET-DNA into smaller “digestible” fragments.

Next, we explored the mechanistic basis of defective DNase secretion in 7-KC-treated macrophages. Since, atherosclerotic plaque macrophages and 7-KC-treated foamy macrophages are known to be in a state of chronic compensated ER stress^28,29^ and ER stress is known to affect secretory processes^30^, we tested whether the defective DNase secretion during hypercholesterolemia could be mediated by ER stress. Consistent with previous studies, we observed that 7-KC-treated macrophages were under ER stress as demonstrated by upregulation of the spliced form of XBP1 (**Supplementary Figure 5C**). To relieve ER stress, we incubated 7-KC-treated macrophages with tauroursodeoxycholic acid (TUDCA), a chemical chaperone that is known to relieve ER stress both *in-vitro* and *in-vivo*^31^. Interestingly, we observed that TUDCA-mediated ER stress relief (**Supplementary Figure 5C**) reversed the impairment in DNA-induced DNase secretion in 7-KC-treated macrophages (**Figure 7G**). Consistent with an increase in secreted DNase activity, the efficiency of extracellular DNA clearance and NET-DNA engulfment of TUDCA-treated 7KC-macrophages was restored to a level similar to that of control macrophages (**Figure 7H, 7I, and Supplementary Figure 5D**). Similar results were obtained when 7-KC-treated macrophages were incubated with azoramide^32^ (**Supplementary Figure 5E**), another known ER stress reliever, which suggests that the observed phenotype is primarily due to reversal of ER stress and unlikely to be an off-target effect of the specific reagents used. These data taken together suggest that ER stress may be a causal factor leading to the impairment in DNA-induced DNase response during atherogenic dyslipidemia.

### ER stress relief rescues the defective DNA-induced DNase response in hypercholesterolemic mice

Based on the *in-vitro* data above, we tested whether relieving ER stress *in-vivo* by administration of TUDCA could rescue the defective DNA-induced DNase response in hypercholesterolemic mice. Towards this end, we induced severe hypercholesterolemia in *Apoe*^*−/−*^ mice by feeding them western-type diet for 3 weeks. During this period, one group of mice was administered TUDCA (150 mg/kg intraperitoneally, daily)^33^ while the control group was administered PBS. As expected, TUDCA-treated mice had significantly lower ER stress as demonstrated by a decrease in the levels of spliced form of XBP1 (**Figure 8A**). It is important to note that body weight and metabolic parameters including blood glucose, plasma cholesterol, and triglyceride were similar between TUDCA-treated mice and control mice (**Supplementary Figure 6A-D**). Interestingly, we observed that the basal level of extracellular DNA was lower in the TUDCA-treated mice as compared with control mice (**Figure 8B**). Consistent with the lowering of serum extracellular DNA levels, TUDCA-treated mice showed elevated basal serum DNase activity (**Figure 8C**). Next, we injected NET-DNA in these mice to analyze the DNA-induced DNase response and efficiency of DNA clearance. We observed that the TUDCA-treated mice showed a significantly higher total plasma DNase activity as compared with control mice upon injection of NET-DNA (**Figure 8D**). DPZ revealed that this increase in plasma DNase activity was contributed by an increase in the levels of both DNase1 and DNase1L3 (**Figure 8E**). Most importantly, the rate of clearance of extracellular DNA as measured by the resolution interval was significantly shortened in TUDCA-treated mice as compared with control mice (**Figure 8F and G**). Consistent with the accelerated clearance of extracellular DNA, the expression of pro-inflammatory genes in the spleen (**Supplementary Figure 6E**) and the expression levels of pro-inflammatory cytokines in the plasma was significantly decreased in the TUDCA-treated mice (**Supplementary Figure 6F**). These data suggest that atherogenic dyslipidemia-induced ER stress mediates the defective DNA-induced DNase response, leading to impaired extracellular DNA clearance and exacerbated systemic inflammation.

**Figure 8.**
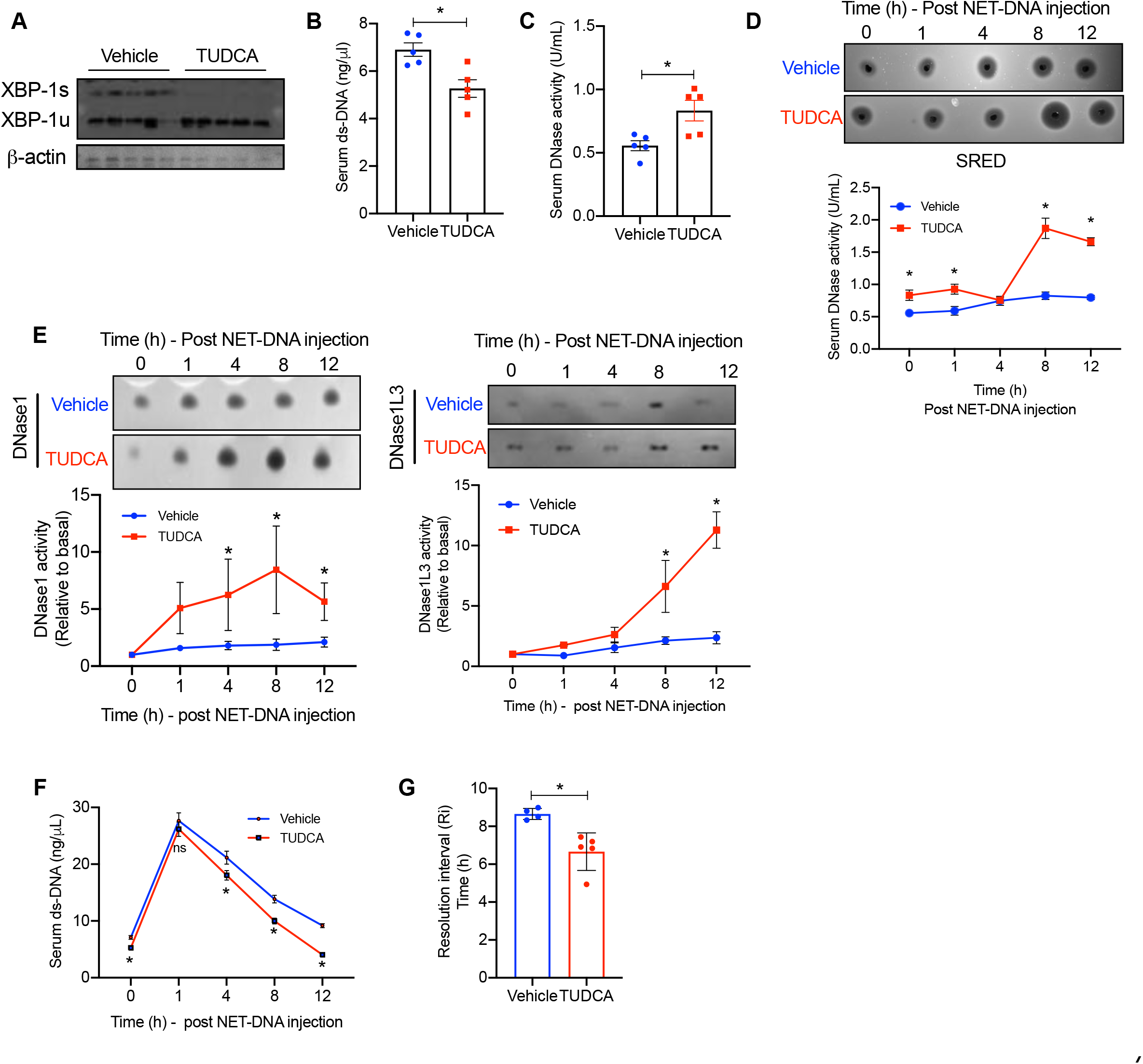
TUDCA rescues the defective DNA-induced DNase response in hypercholesterolemic *Apoe*^*−/−*^ mice. 10-wk old *Apoe*^*−/−*^ mice were fed WD for 3 wks with concomitant daily administration of either vehicle or TUDCA intraperitoneally. (A) Immunoblot analysis of the expression levels of unspliced and spliced forms of XBP-1 in whole cell lysates from peritoneal macrophages of vehicle and TUDCA-treated mice. -actin was used as loading control. (B) Analysis of basal serum extracellular ds-DNA levels and (C) total DNase activity in vehicle and TUDCA-treated mice. (D) SRED-based quantification of total DNase activity, (E) DPZ-based analysis of Dnase1 and DNase1L3 expression levels, and (F) measurement of serum extracellular ds-DNA levels at indicated time points post-injection of NET-DNA in vehicle and TUDCA-treated mice. (G) Quantification of resolution interval in vehicle and TUDCA-treated mice. n = 5 mice per group. *, p < 0.05; ns, no significant difference.

## Discussion

There is overwhelming evidence of the pathogenic role of excessive NETosis and persistence of extracellular DNA in promoting disease progression with consequent adverse pathological and clinical outcomes^2,34^. Previous studies have highlighted the critical role of plasma DNases in the clearance of extracellular DNA and maintenance of tissue homeostasis^4^. Indeed, lower basal DNase activity, either due to mutations in DNase1 or DNase1L3^35,36^, or the presence of plasma DNase inhibitors^37,38^, has been implicated to adversely impact disease outcomes, particularly in autoimmune lupus, myocardial infarction, sepsis, acute pancreatitis, cancers, NAFLD, and cirrhosis^4^. In this context, our data provides evidence for a role for hypercholesterolemia-induced dysregulation of DNA-induced DNase response as a critical additional mechanism for sub-optimal DNase levels and impaired extracellular DNA clearance.

Previous studies have demonstrated a correlation between an increase in extracellular DNA levels with concomitant increase in plasma DNase activity such as in myocardial infarction, sepsis, and acute pancreatitis^12,14^. Since, liver is one of the major source of DNase1^39^, it was not clear whether in these clinical conditions of acute inflammation, the increase in DNase activity was part of an acute phase reaction or a specific response to an increase in plasma extracellular DNA. Our study demonstrates that an increase in extracellular DNA levels can be sensed by certain tissue or cell types and set in motion a response by secreting DNases to promote degradative clearance of DNA and restoration of tissue homeostasis.

While it is evident that NET-DNA is sensed by certain cellular receptors which then mediates the DNase secretion response, the nature and identity of this receptor(s) is currently unknown. Since the increase in DNase secretion *in-vivo* and *in-vitro* is elicited only upon exposure to NET-DNA and not by nuclear DNA stripped off its proteins, our data suggests that the DNase response is mediated by sensing the DNA-protein complex. In this context, a recent study identified CCDC25 as a plasma membrane localized receptor on cancer cells that binds exclusively to the DNA component of NET-DNA and mediates downstream signaling involving activation of ILK-β-parvin pathway that enhances cell motility and cancer metastasis^40^. Given that the DNase response we observe is elicited only by the DNA-protein complex and not by the naked-DNA, it is unlikely that CCDC25 is involved in this response. This raises the interesting possibility of the existence of additional NET-DNA binding receptors which might be expressed in a tissue or cell type specific manner that elicits functionally distinct responses.

Additionally, in contrast to the current understanding that myeloid cells such as macrophages and dendritic cells are the major source of DNase1L3^21^ while DNase1 is reportedly secreted primarily by non-hematopoietic cells, we demonstrate for the first time that both murine and human macrophages exposed to extracellular DNA have the ability to secrete significant quantities of DNase1 that is functionally relevant for DNA degradation *in-vitro*. In addition to the extracellular degradation of DNA by macrophage-secreted DNase, our data also suggests an important role for the secreted DNases in facilitating macrophage-mediated engulfment via digestion of DNA into smaller fragments, which leads to enhanced intracellular phago-lysosomal degradation of DNA. These data are consistent with a previous study that demonstrated that EDTA, which chelates divalent cations such as Ca^2+^, Mg^2+^, and, Mn^2+^ and blocks DNase activity, leads to decreased clearance of NET-DNA by macrophages. Since, EDTA could have several non-specific effects including inhibitory activity on phagocytic processes, our approach of specifically inhibiting the activity of DNase1 and DNase1L3 using aprotinin and plasmin is a more definitive demonstration of the co-operative role played by macrophage-secreted DNase in phagocytic clearance of NET-DNA. Future studies using mice with macrophage-specific knockout of DNase1 and DNase1L3 (DNase1 ^flox/flox^ / DNase1L3^flox/flox^ mice are currently unavailable) will be required to specifically understand the role of locally produced macrophage secreted DNases in extracellular DNA clearance and inflammation resolution *in-vivo*.

Since persistent extracellular DNA is highly pro-inflammatory, its prompt clearance is essential for maintaining immune homeostasis. Consistent with this notion, we observe that the DNA-induced DNase response is a rapid response demonstrating significant elevation in DNase levels within 4 hours of elevation of extracellular DNA. Interestingly, our study demonstrates that this response is not mediated by either a transcriptional or translational upregulation of DNases. Since it is known that DNases are localized to vesicles within the cytoplasm of cells^41^, it raises the interesting possibility that the rapid response may be mediated by quick release of DNases by exocytosis from pre-formed granules similar to that reported in exocrine pancreas.

A previous study^19^ demonstrated that exogenous administration of DNase1 decreases atherosclerotic lesion area in *Apoe*^*−/−*^ mice fed a high-fat chow diet for 6 weeks suggesting a beneficial effect of DNase1-mediated clearance of NET-DNA in lowering inflammation and protecting against early atherosclerosis. Similarly, the administration of DNase1 improved the regression of atherosclerotic plaques in diabetic *Ldlr*^*−/−*^ mice when switched to a chow diet^20^. Complementing the findings of these studies, our results in a mouse model of advanced atherosclerosis, demonstrate that restoration of the defective sub-optimal DNase response during hypercholesterolemia by exogenous administration of DNase1 aids in the lowering of local and systemic inflammation, decreases necrotic area in the plaque, and increases lesional collagen content. It is important to note that large areas of plaque necrosis and decreased collagen content are indicative of “rupture-prone” vulnerable plaque in humans^42^, which are causally associated with the development of clinical outcomes including myocardial infarction and stroke. In this context, out data suggest that DNase1 administration could have beneficial effects even in established advanced atherosclerotic plaques resulting in significant atherosclerotic plaque remodeling towards a stable plaque phenotype. Since, DNase1 is approved for clinical use in patients with cystic fibrosis^43^, further studies to test the efficacy of DNase1 as a therapeutic target in high-risk patients with advanced atherosclerosis is warranted.

Using 7-ketocholesterol, a pathophysiologically relevant ER stressor in the context of atherogenic dyslipidemia, we have demonstrated that ER stress impairs the DNA-induced DNase response and results in delayed clearance of extracellular DNA. In theory, ER stress could affect the recognition of extracellular DNA by decreasing the expression level of the putative NET-DNA receptor thereby decreasing the efficiency of DNA sensing. However, this seems unlikely, since we did not observe significant difference in the NET-DNA binding efficiency of ER stressed-macrophages (unpublished data). Alternately, ER stress could affect the signaling downstream of the putative NET-DNA receptor and impair trafficking of DNase-containing vesicles to the plasma membrane. Indeed, ER stress is known to impair secretory processes by inhibiting focal exocytosis in an ATF4-TRB3 dependent manner in pancreatic β-cells during diabetes^44^. Whether such a mechanism is responsible for the decreased secretion of DNases by macrophages and other cell types remains to be explored.

Interestingly, our demonstration that ER stress leads to the maladaptive DNase response raises the interesting possibility that this phenomenon may be broadly applicable to several diseases associated with chronic ER stress such as diabetes, COPD, NAFLD etc. Although this is a speculation and needs a separate study, consistent with this concept, increased levels of plasma extracellular DNA is considered an independent risk factor and an indicator of poor prognosis in NAFLD^45^ and COPD^46^.

In summary, our study demonstrates the existence of a DNA-induced DNase response which acts as a critical feedback mechanism to maintain extracellular DNA levels within narrow physiological limits. Further, we suggest that an impairment of this DNA-induced DNase response, such as during hypercholesterolemia, impairs inflammation resolution and promotes disease progression with consequent adverse clinical outcomes. These findings open novel therapeutic opportunities to explore strategies aimed at reversing the defective DNA-induced DNase response during hypercholesterolemia in an effort to stall or reverse the progression of atherosclerosis and other chronic inflammatory diseases.

## Materials and Methods

### Animals and animal maintenance

C57BL6 mice were bred at the animal facility of CSIR-Institute of Genomics and Integrative Biology, New Delhi. *Apoe*^*−/−*^ mice on a C57BL/6 genetic background was obtained from CSIR-Centre for Cellular and Molecular Biology, Hyderabad, India, through a material transfer agreement and were bred as homozygotes and maintained at the animal facility of CSIR-Institute of Genomics and Integrative Biology. To induce short-term hypercholesterolemia, 8-10 wk old male or female *Apoe*^*−/−*^ mice were fed a western-type diet (D12079B, Research Diets, 40% Kcal from fat, 0.2% cholesterol) ab-libitum. To generate advanced atherosclerotic plaques, 8-10 wk old male or female *Apoe*^*−/−*^ mice were fed a Western-type diet ab-libitum for 16 wks. All animal protocols used in this study were approved by the Institutional Animal Ethics Committee (IAEC) of CSIR-Institute of Genomics and Integrative Biology, New Delhi.

### Human plasma isolation

5 mL of peripheral blood was collected by venipuncture into vacutainer (BD) tubes from healthy adults. The samples were centrifuged at 2000 × g for 15 minutes to harvest the plasma. The isolated plasma was aliquoted and stored at −80 °C until further use. All studies involving human volunteers were conducted after obtaining informed consent and with the approval of the Institutional Human Ethics Committee of CSIR-Institute of Genomics and Integrative Biology, New Delhi.

### Human peripheral blood mononuclear cell isolation and macrophage differentiation

Human peripheral blood mononuclear cells (PBMC) were isolated as described previously. Briefly, 1:2 diluted blood was layered on Histopaque-1077 (Sigma) and centrifuged at 400 × g for 30 min in a swing-bucket rotor without brake. The supernatant was discarded and the PBMCs were isolated from the interface and were cultured in DMEM supplemented with 10% (vol/vol) heat-inactivated FBS, 10 U/mL penicillin, 100 mg/mL streptomycin. Post 6 h of seeding, fresh complete media supplemented with 20 ng/mL human M-CSF was added. The spent media was replaced on day 3 followed by culture of cells for upto 7 days to allow macrophage differentiation.

### Cell Culture

Human HL-60 and THP-1 cells (obtained from National Centre for Cell Science, Pune, India) were cultured in RPMI-1640 supplemented with 1 mM sodium pyruvate, 10% fetal bovine serum (FBS), 1000 U/mL penicillin and 10000 μg/mL streptomycin and 25μg/mL of amphotericin B while mouse RAW264.7 and L-929 cells were cultured in DMEM (high-glucose) supplemented with 10% FBS, 1000 U/mL penicillin and 10000 μg/mL streptomycin and 25μg/mL amphotericin B. All cells were cultured at 37°C in a humidified 5% CO_2_ incubator. THP-1 cells were seeded at 0.5×10^6^ cells/well in a 12-well plate in complete media supplemented with 100 nM PMA to differentiate them in to macrophage-like cells. To collect GMCSF enriched media, L-929 cells were seeded in 75 cm^2^ cell culture flask (Corning®) and after 90% confluency of the cells, the media was replaced and cells were grown for 7 days and the cell culture supernatant was collected. The collected supernatant was centrifuged at 1500× g for 5 minutes followed by filtration through a 0.22 μm filter and stored at −80 °C until use.

### Bone marrow-derived macrophage culture

Bone marrow cells from 8-12 weeks old male or female mice were flushed using a syringe fitted with a 26G needle followed by passing through a 100 μm cell strainer to eliminate cell clumps and debris. Cells were pelleted by centrifuging at 1200× g followed by RBC lysis using commercially available RBC lysis buffer (Sigma, R7757). The cells were then pelleted and resuspended in DMEM supplemented with 10% (vol/vol) heat-inactivated FBS, 10 U/mL penicillin, 100 mg/mL streptomycin, and 20% (vol/vol) L-929 cell culture supernatant for 7 days to allow macrophage differentiation.

### Isolation of NET-DNA and systemic injection

HL-60 cells cultured in RPMI supplemented with 10% FBS were incubated with 1 μM All-trans retinoic acid (ATRA R2625-50MG, Sigma) for 4 days to induce neutrophil-like differentiation following which they were treated with 100 nM PMA for 4 h to induce NETosis^15^ and release of NETs. The plate-adherent NET-DNA was scraped and collected along with the culture supernatant. Cells and cell debris were pelleted by low-speed centrifugation at 450 x g for 10 min. The NET-rich supernatant thus obtained was further centrifuged for 10 minutes at 18,000 × g at 4 °C to pellet NET-DNA which was then resuspended in ice-cold PBS. DNA concentration was measured in the sample obtained using spectroscopy absorbance at 260/280 nm. The isolated NET-DNA was diluted in 1X-PBS to a final concentration of 40 μg in 300 μl and injected intravenously via the lateral tail vein of 10 wk-old C57BL6 mice. 20 μl of blood was collected by tail bleed at periodic intervals of 1, 4, and 8 h post-injection for analysis of extracellular DNA and DNase activity. The mice were euthanized at 12 h post-injection of NET-DNA by an overdose of thiopentone sodium followed by intracardiac puncture to isolate blood for detailed analysis of several biochemical and inflammatory parameters.

### Measurement of plasma and serum extracellular ds-DNA

EC-DNA concentration was measured in the plasma or serum sample using Qubit™ dsDNA HS Assay Kit using according to the manufacturer’s protocol (Thermofisher Cat#Q32851)

### Quantification of total DNase activity by single radial enzyme diffusion (SRED) assay

SRED for measuring total DNase activity in the plasma, serum, or cell culture supernatant was conducted as described previously^3^. Briefly, 55 μg/mL salmon testes DNA was resuspended in a buffer containing Mn^2+^ (20 mM Tris-HCl pH 7.8, 10 mM MnCl_2_, 2 mM CaCl_2_, and 2X ethidium bromide). The DNA solution was heated at 50°C for 10 minutes and mixed with an equal volume of 2% agarose (Sigma-Aldrich). The mixture was poured onto a plastic casting tray and left at room temperature till it solidified. Wells of approximately 0.1 mm size were created using a 20 μL pipette tip. 2 μl of murine serum or 4 μl of human plasma were loaded into wells and gels were incubated for 6 hours at 37°C in a humidified chamber followed by image acquisition using UV excitation in Gel Doc system (Syngene G: BOX XX6). The area of DNA degradation was analyzed on Fiji (ImageJ) and the DNase activity in the sample was calculated by interpolation from standards of known DNase activity.

### Denaturing polyacrylamide gel electrophoresis zymography (DPZ) for measurement of DNase1 and DNase1L3 activity

DPZ was performed as described previously with some modifications^3,16^. Briefly, sodium dodecyl sulphate (SDS)—polyacrylamide gels were prepared with 4% (v/v) stacking gels without DNA and 10% (v/v) resolving gels containing 200 μg/mL of salmon testes DNA (Sigma-Aldrich, Germany). For the detection of DNase1, 0.5 μl of murine serum or 5 μl of cell culture supernatant was mixed with 12 μl of water and 5 μl SDS gel-loading buffer. The mixture was loaded onto wells and electrophoresis was carried out at 120 V in a Tris-glycine electrophoresis buffer (25 mM Tris, 192 mM glycine, 0.1% (w/v) SDS, pH 8.7). After electrophoresis, SDS was removed by washing the gels twice for 30 min each at 50°C with 10 mM Tris-HCl pH 7.8 following which the proteins were refolded by incubating the gels overnight at 37°C in a re-folding buffer (10 mM Tris-HCl pH 7.8, 3 mM CaCl_2_, 3 mM MgCl_2_). Images were acquired with UV excitation in a Gel Doc system followed by image quantification using Fiji.

For the detection of DNase1L3, 2 μl of serum was mixed with 12 μl of water, 5 μl SDS gel-loading buffer and 1 μl of beta-mercaptoethanol (β-ME) which inactivates DNase1 via reduction of its disulphide bridges. The mixture was heated for 5 minutes and loaded onto the gels. Electrophoresis and subsequent washing were carried out as described for DNase1. The proteins were refolded by sequential incubation of the gel for 48 h at 37°C in refolding buffer - 1 (10 mM Tris-HCl pH 7.8 containing 1 mM β-ME) followed by a further 48 h incubation at 37°C in refolding buffer - 2 (refolding buffer 1 supplemented with 3 mM CaCl_2_ and 3 mM MnCl_2_). Images were acquired with UV excitation in a Gel Doc system followed by image quantification using Fiji.

### Gene expression analysis by qPCR

RNA was isolated using RNeasy Mini Kit (QIAGEN) and reverse transcribed using a first-strand cDNA synthesis kit (PrimeScript™ 1st strand cDNA Synthesis Kit, Takara) according to the manufacturer’s protocol. Real time-PCR was conducted by SYBR green chemistry on a Roche LightCycler 480. The details of primers used in this study are provided below.

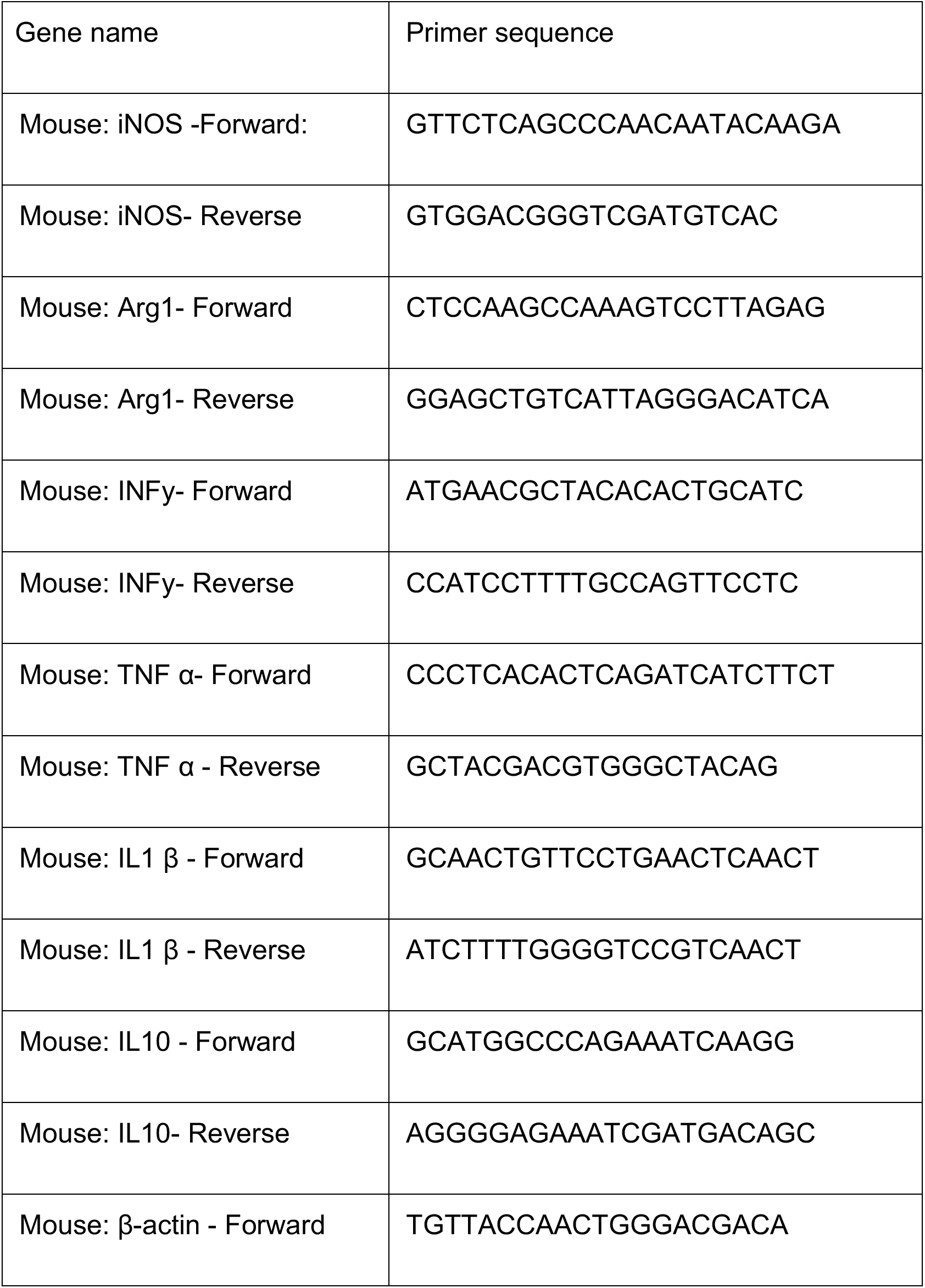
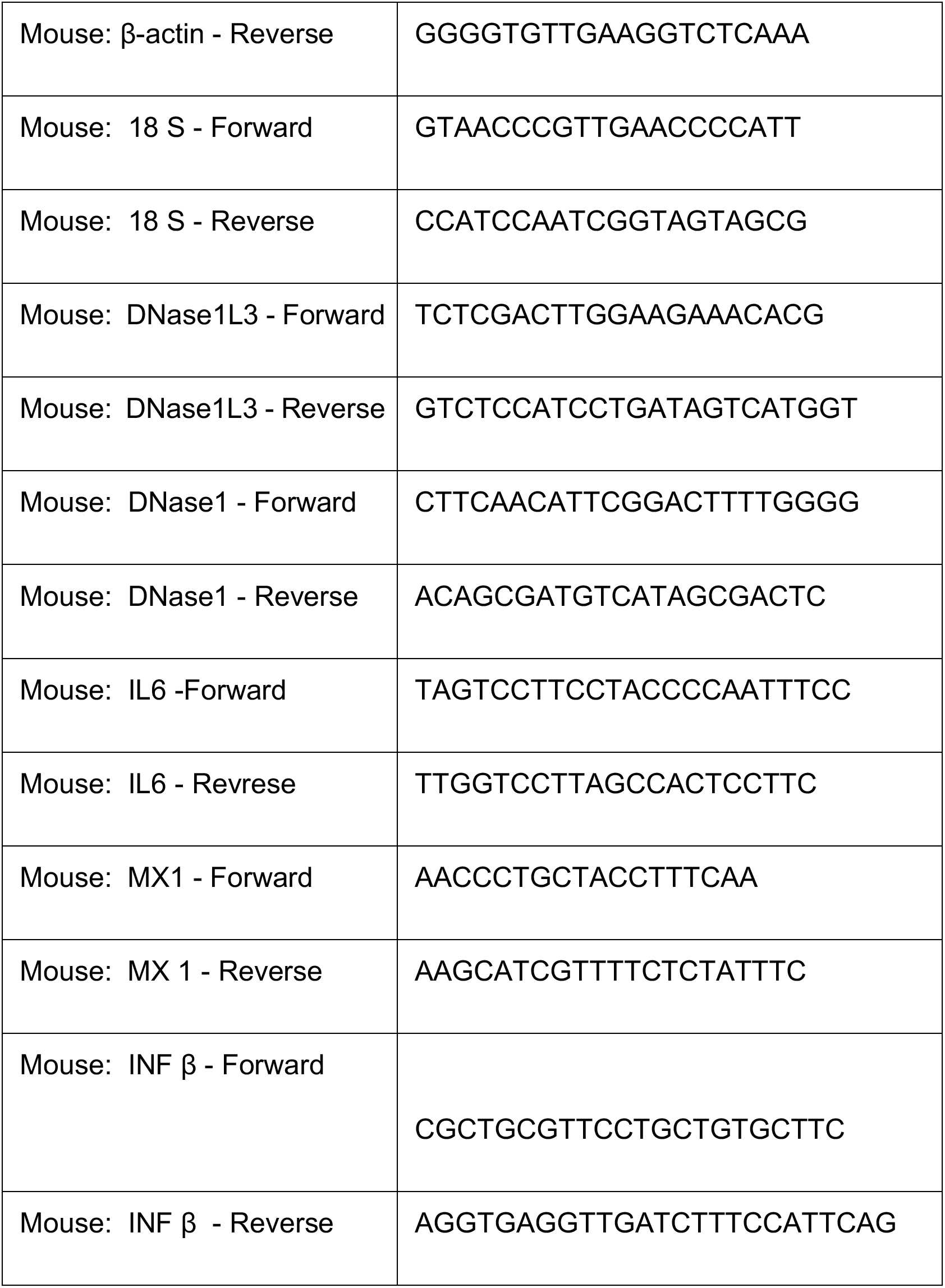

### Analysis of NET-DNA engulfment and clearance

BMDMs or THP-1 cells were seeded at 0.5×10^6^ cells/well in a 12-well plate and incubated overnight. For NET-DNA engulfment assay, the spent media was removed following which the cells were washed with 1X-PBS and incubated with 10 μg Sytox green-labeled NET-DNA for 2 h. Post-incubation, unengulfed NET-DNA were washed off and images were acquired on a fluorescence microscope. The efficiency of engulfment was quantified as percent macrophages that showed intracellular NET-DNA fluorescence. For the purpose of measuring total NET-DNA clearance efficiency, cells were seeded as above and incubated with 10 μg NET-DNA for 12 h following which the remaining quantity of NET-DNA in the supernatant was analyzed by measuring absorbance using Qubit.

### Immunofluorescence staining

Coverslips were coated with poly-L-lysine for 1 h prior to seeding HL-60 cells at a density of 0.5×10^6^ and incubated overnight. Spent media was removed gently and the cells were incubated with 100 nM PMA for 4 h following which the cells were fixed with 4% paraformaldehyde. Samples were blocked with 3% FBS in 1X-PBS followed by incubation with anti-CitH3 antibody (1:100, NB100-571350, Novus Biologicals) and Alexa fluor 488-conjugated anti-rabbit antibody (1:1000, Thermo Fisher Scientific). The nuclei were counterstained using DAPI (1:10000, Sigma-Aldrich). Images were acquired on a fluorescence microscope.

### Immunoblotting

Cells were lysed in 2X-Laemmli sample buffer containing 1mM β-mercaptoethanol and heated at 100°C for 10 min. The samples were loaded onto 10% SDS-polyacrylamide gel and electrophoresis was carried out at 120 V using Tris-glycine buffer. Western transfer on to 0.45 μm PVDF membrane was conducted at 35A for overnight. The membrane was blocked using 5% skim milk and the primary antibody (1:1000) was added in 1% skim milk and incubated for three hours at room temperature. Three washes were given with 0.1% PBST for 10 mins each. Secondary antibody (1:5000) in 1% skim milk was added and incubated for 1 h at room temperature. Three washes were given with 0.1% PBST for 10 mins followed by incubation with enhanced chemiluminescence substrate and image capture in a gel documentation system.

### *In-vitro* DNA degradation assay

*In-vitro* digestion was performed using 2 μL of murine serum mixed with 5 μg of input DNA and incubated at 37°C for 2.5 h. To examine the DNase activity in cell culture supernatants, 1mL of supernatant was concentrated up to 100-fold using a 10 KDa centricon. A reaction mixture was made by mixing 10 μL of the concentrate with 5 μg of input DNA and DNase buffer (10mM tris-HCl, 3mM CaCl_2_, 3mM MnCl_2_) and incubated at 37°C for 12 h. The reaction mixture was then loaded on to agarose gel and DNA degradation was analyzed following electrophoresis.

### Generation of necrotic and apoptotic cells

Briefly, to generate apoptotic cells, HL-60 cells were irradiated under a shortwave UV lamp for 7 minutes and incubated under normal cell culture conditions for 2-3 hr. To induce necrosis, HL-60 cell suspension was incubated at 56°C for 15 minutes.

### Multiplex ELISA

A panel of inflammatory cytokines were assayed using commercially available ProcartaPlex (12-plex) immunoassays (Invitrogen). 25 μL of serum was used for analysis as per manufacturer’s protocol in a bead compatible assay analyzer (Magpix System, Luminex Corporation).

### Metabolic profiling

Blood glucose was measured in 5h-fasted mice using commercially available glucose strips and glucometer (Accucheck). For the measurement of plasma total cholesterol and triglycerides, the mice were fasted overnight followed by collection of 20 μL of blood from the lateral tail vein. Total cholesterol and triglyceride were quantified using Cholesterol E kit and Triglyceride M Color B kit from Wako as per manufacturer’s instructions.

### Aortic root atherosclerotic lesion analysis

Eight-to-ten-weeks-old *Apoe*^*−/−*^ female mice were fed a western-type Diet for 16 weeks. Immediately after euthanasia, the mice were perfused with 1X-PBS by cardiac puncture into the left ventricle, and aortic roots, along with heart were collected and fixed in 10% neutral buffered formalin. Later, the tissues were embedded in paraffin and 8 μm-thick sections were cut using a microtome. 50-serial sections starting from the first appearance of the aortic valve were collected for analysis of aortic root atherosclerosis. Total lesion area and necrotic area were analyzed by H&E staining of 6 sections per mouse spanning the entire aortic root. For analysis of lesional collagen, the sections were stained with Mason Trichrome stain (Sigma-Aldrich) as per manufacturer’s protocol.

### Immunohistochemistry

Paraffin-embedded tissue specimens were sectioned, de-paraffinized with xylene, and rehydrated in decreasing concentrations of ethanol followed by 3 washes with 1X-PBS. Antigen retrieval was performed in boiling Sodium citrate buffer (10 mM Sodium citrate, 0.05% Tween-20, pH 6.0) or using Proteinase-k (20 μg/mL). The sections were then blocked with 3% fetal bovine serum in 1X-PBS for 30 min, followed by overnight - incubation at 4°C with anti-F4/80 (1:50), anti-CitH3 (1:50), anti-Dnase1 (1:50), anti-Dnase-1L3 (1:50), anti-Sm-actin (1:50). After 3 washes with 1X-PBS, the sections were further incubated with fluorescently-labeled secondary antibodies and counterstained with DAPI. Images of the stained sections were captured using a fluorescence microscope (Nikon Eclipse, Ti) and image analysis was performed using Fiji (ImageJ).

## Acknowledgements

This study was supported by research funding from the DBT-Wellcome Trust India Alliance Intermediate Fellowship (MS) and Barts Charity, UK (MS). UD and PB were supported by Senior research fellowship from the Department of Biotechnology, India, and Council of Scientific and Industrial Research, India, respectively. We thank Dr. Shantanu Sengupta and Dr. Vivek Natarajan (CSIR-Institute of Genomics and Integrative Biology, New Delhi) for helpful discussions throughout the project.

## Author contributions

UD, PB, SN, and VP designed and conducted experiments and analysed the data. AA conducted analysis of publicly available microarray and RNA-sequencing data. UD and MS conceptualized the study, wrote, and edited the manuscript.

