## Supplemental Figures for "Hypercholesterolemia Impairs Clearance of Extracellular DNA and Promotes Inflammation and Atherosclerotic Plaque Progression"

#### Supplementary Figure 1

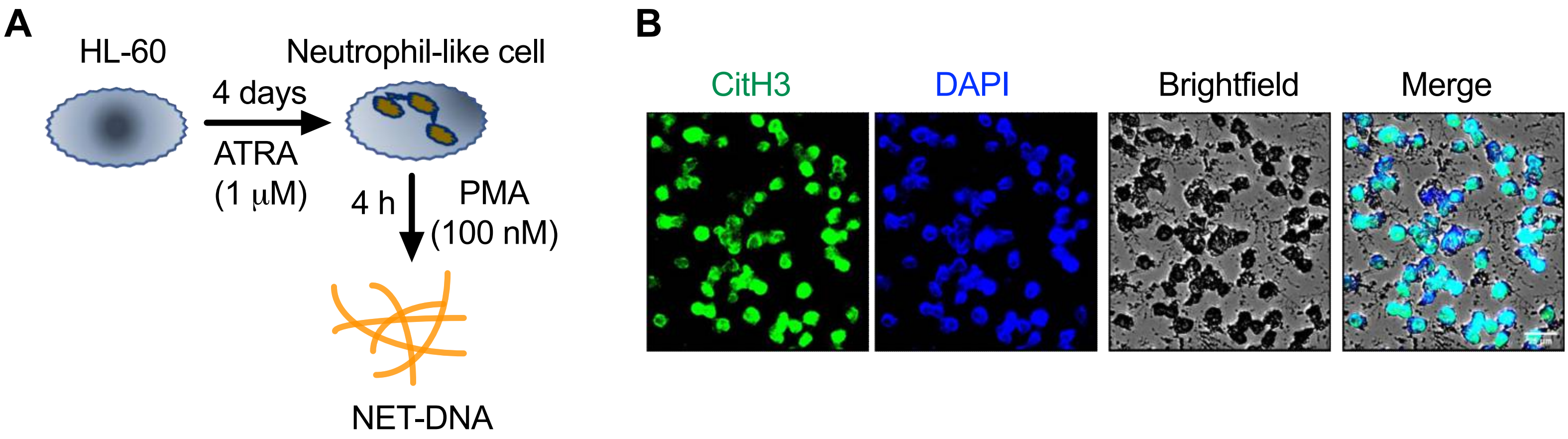


**Supplementary Figure 1. (A)** Schematic depicting the process of differentiation of HL-60 cells into neutrophil-like cells and induction of NETosis using PMA. **(B)** PMA-treated neutrophil-like differentiated HL-60 cells were immunostained with anti-citrullinated H3 antibody (green) and counterstained with DAPI (blue).

### Supplementary Figure 2

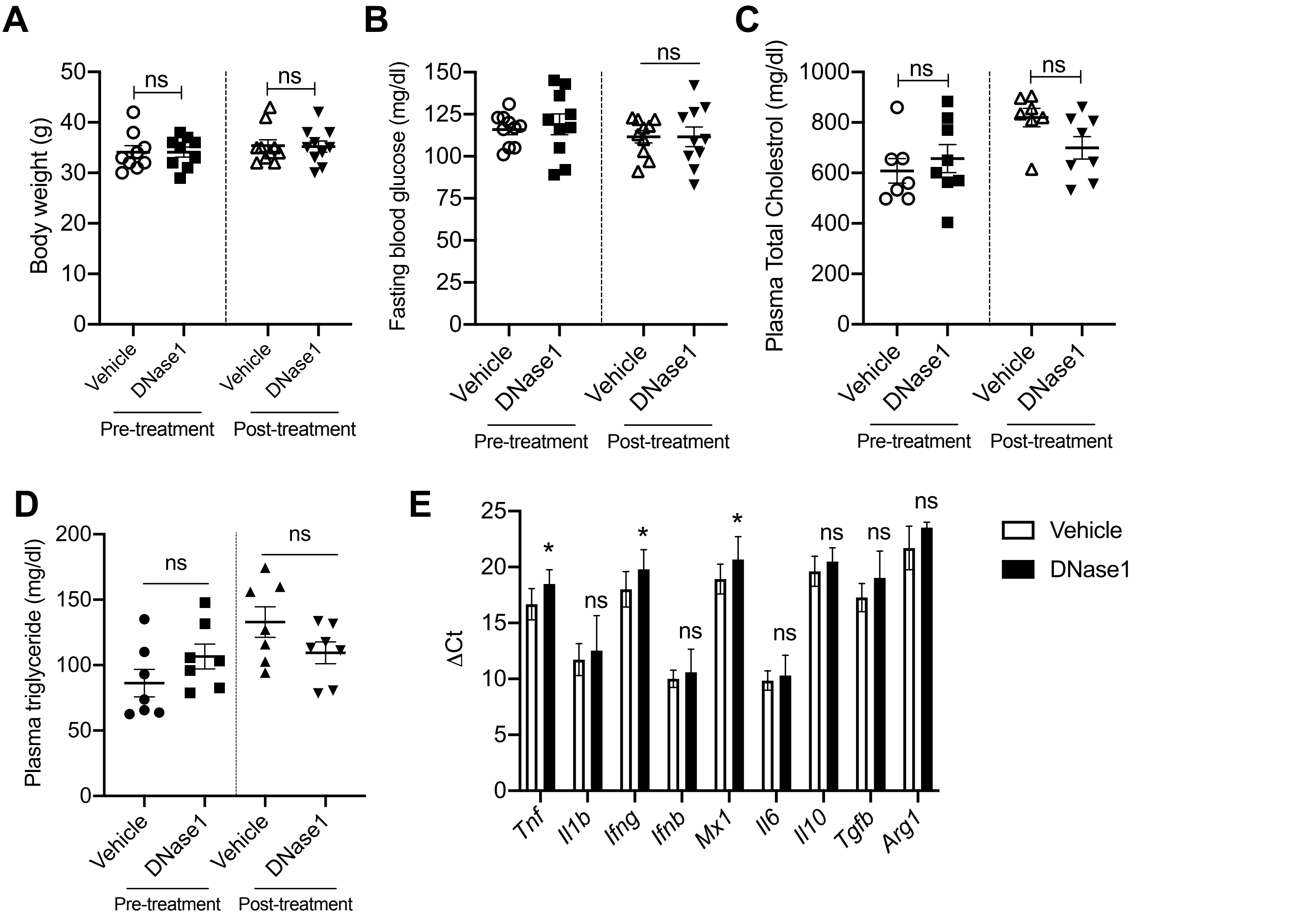


**Supplementary Figure 2.** 16 wk WD-fed *Apoe*$^{-/-}$ mice were administered either vehicle or DNase1 (400 U) intravenously three times a week for 4 wks. **(A)** Body weight, **(B)** blood glucose, **(C)** plasma total cholesterol, and **(D)** plasma triglyceride were quantified prior to and after treatment with either vehicle or DNase. The data represent mean  SEM. n = 10 mice per group. *, $p < 0.05$ using Mann-Whitney test.

### Supplementary Figure 3

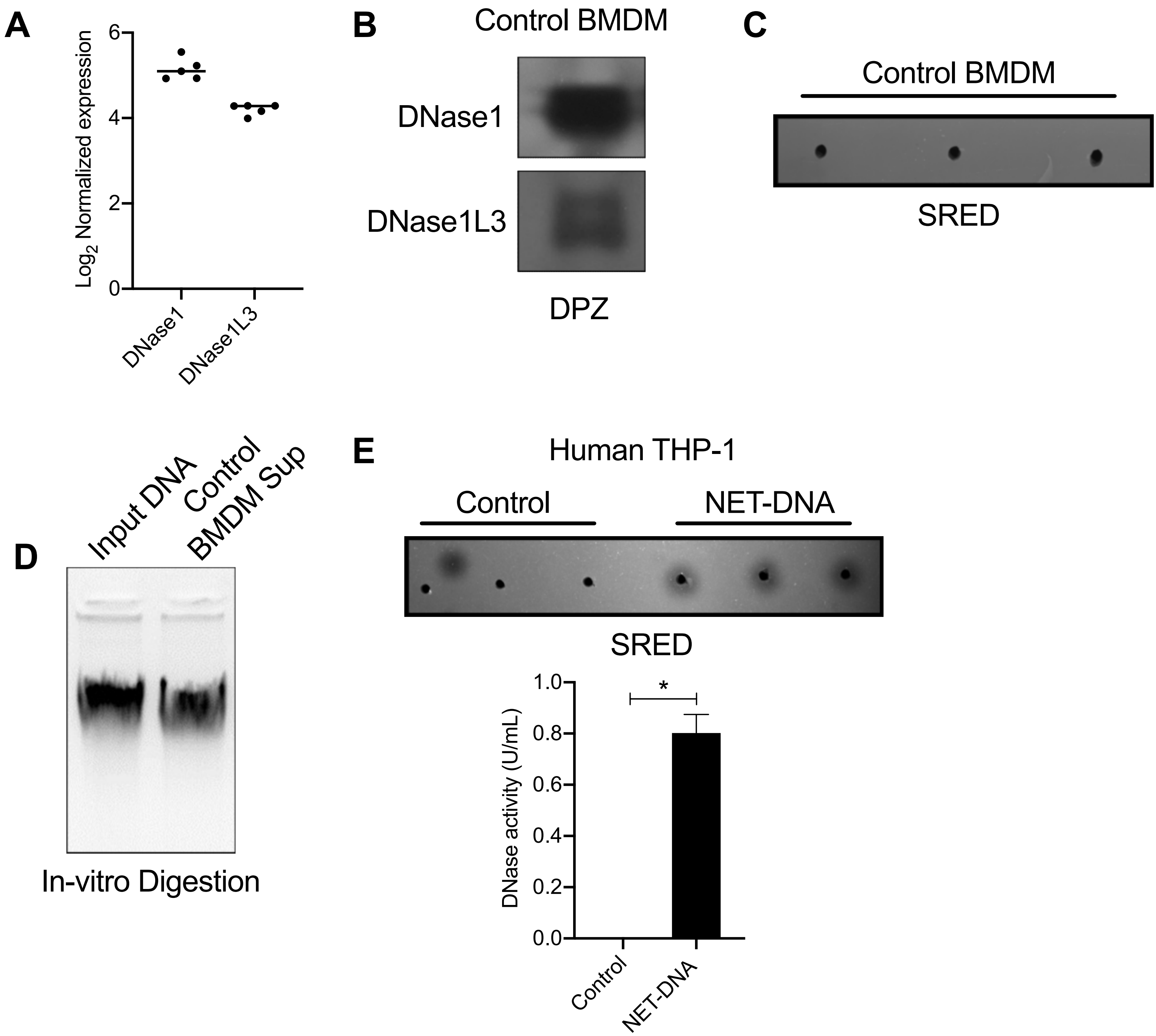


**Supplementary Figure 3. (A)** Raw CEL files were downloaded for the microarray data (GSE70619) and analyzed using Bioconductor (v1.30.1) package Affy (v1.64.0) to get the normalized intensity values (rma normalization). Multiple probes for each gene were collapsed using collapseRows function (default parameters) of package WGCNA (v1.69). **(B)** Depolymerizing gel zymography of 100X concentrated cell culture supernatants from control BMDMs for detection of basal level secretion of DNase1 and DNase1L3. **(C)** SRED for detecting total DNase activity in cell culture supernatant of control BMDM. **(D)** Agarose gel electrophoresis of samples from in-vitro DNA degradation assay (see methods) using cell culture supernatants of control BMDMs. **(E)** SRED-based analysis of total DNase activity in cell culture supernatants of control and NET-DNA-treated human THP-1 macrophages. The bar graph represents mean  SEM. $p < 0.05$ using Mann-Whitney test.

#### Supplementary Figure 4

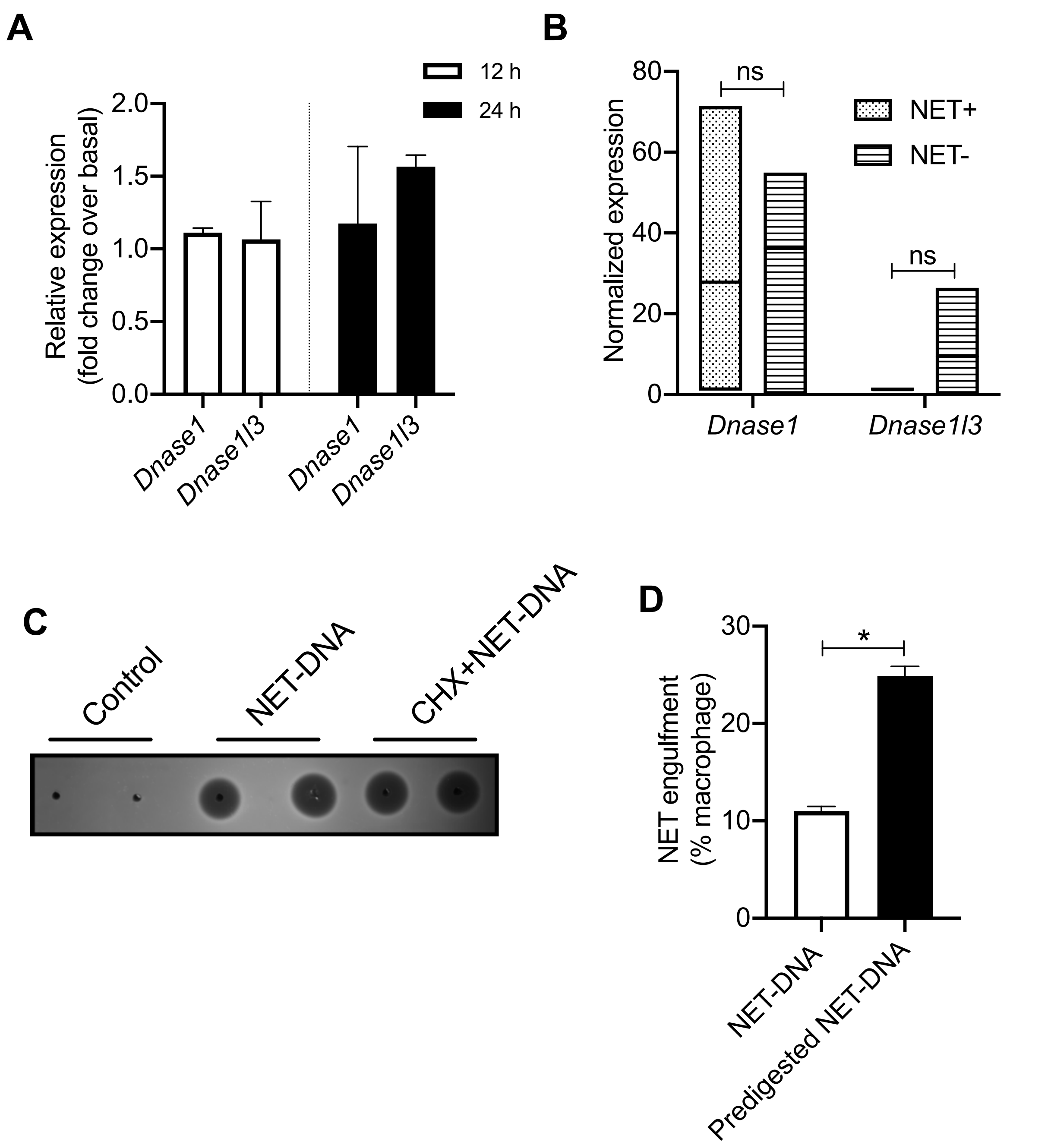


**Supplementary Figure 4. (A)** qPCR-based analysis of mRNA expression of *Dnase1* and *Dnase1l3* in control and NET-DNA-treated macrophages at 12 h and 24 h post-incubation with NET-DNA. **(B)** RNA-sequencing data of plaque macrophages isolated from NET-DNA proximal regions and NET-DNA distal regions were downloaded from GSE145200. The downloaded raw counts data were analyzed using packages splitstackshape (v1.4.8), biomaRt (v2.42.1) and DESeq2 (v1.26.0). Differential expression testing was done using default parameters. **(C)** SRED for analysis of total DNase activity in culture supernatants of control macrophages and macrophages treated with NET-DNA in the absence or presence of cycloheximide (0.5 M). **(D)** Fluorescence microscopy-based quantification of efficiency of engulfment of intact NET-DNA or NET-DNA pre-digested with fetal bovine serum for 1 h at 37C.

#### Supplementary Figure 5

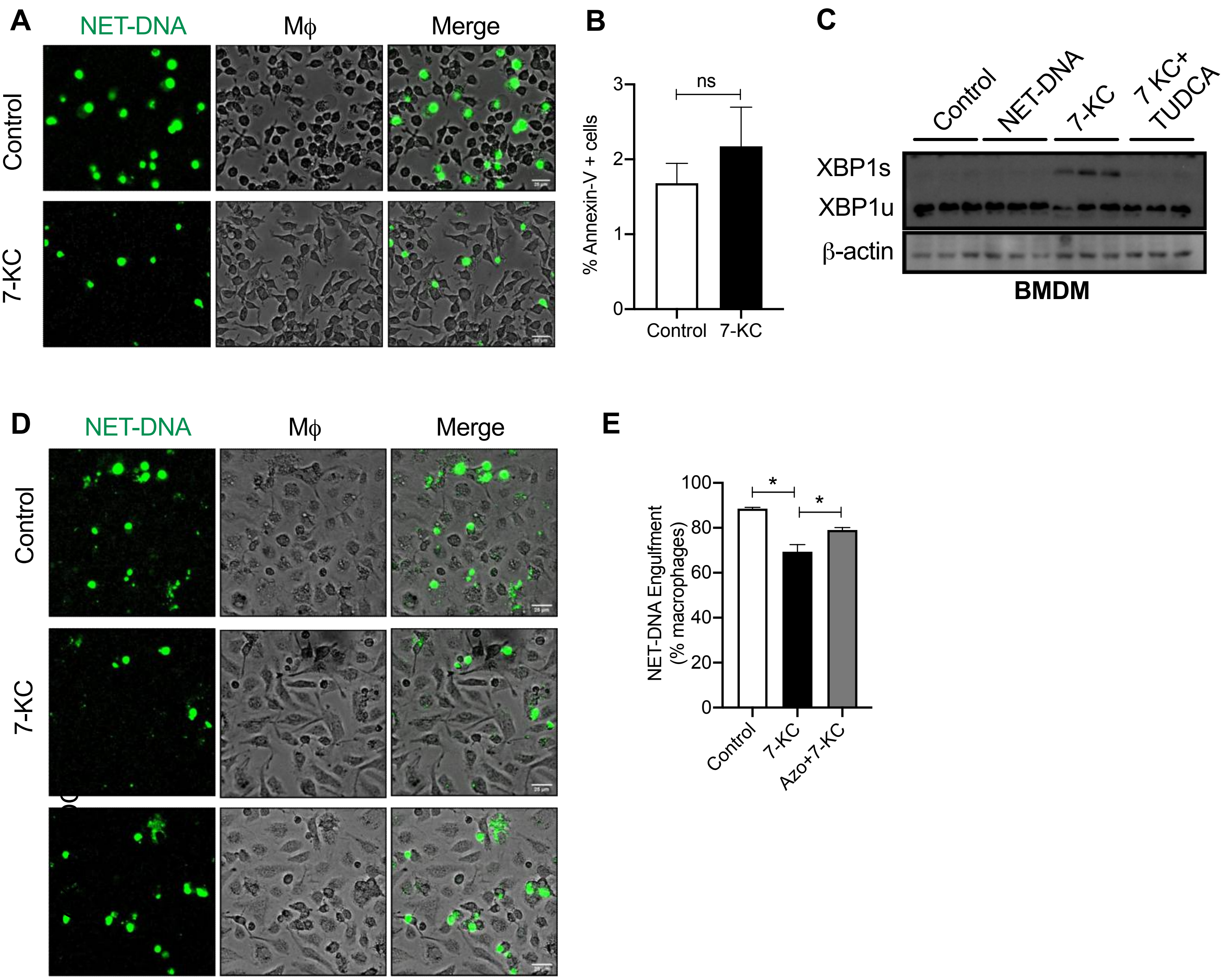


**Supplementary Figure 5. (A)** Representative fluorescence microscopy images used for quantification of efficiency of NET-DNA engulfment. Sytox-green-labeled NET-DNA (green) were incubated with control or 15 M 7-KC-treated macrophages (brightfield). Scale bar, 25 m. **(B)** Mouse BMDMs treated with vehicle or 15 M 7-KC for 18 h were incubated with Annexin-V-Alexa fluor 488 for analysis of annexin-V positive cells by fluorescence microscopy. The bar graph represents mean  SEM derived from three independent experiments. ns, no significant difference as calculated using Mann-Whitney test. **(C)** Control or 15 M 7-KC-treated BMDMs were incubated without or with NET-DNA in the absence or presence of TUDCA as indicated. Immunoblotting for XBP1 was conducted on whole cell lysates after Western transfer. -actin was used as loading control. **(D)** Similar to (A), except that one group of macrophages were additionally co-incubated with Azoramide (10 M).

### Supplementary Figure 6

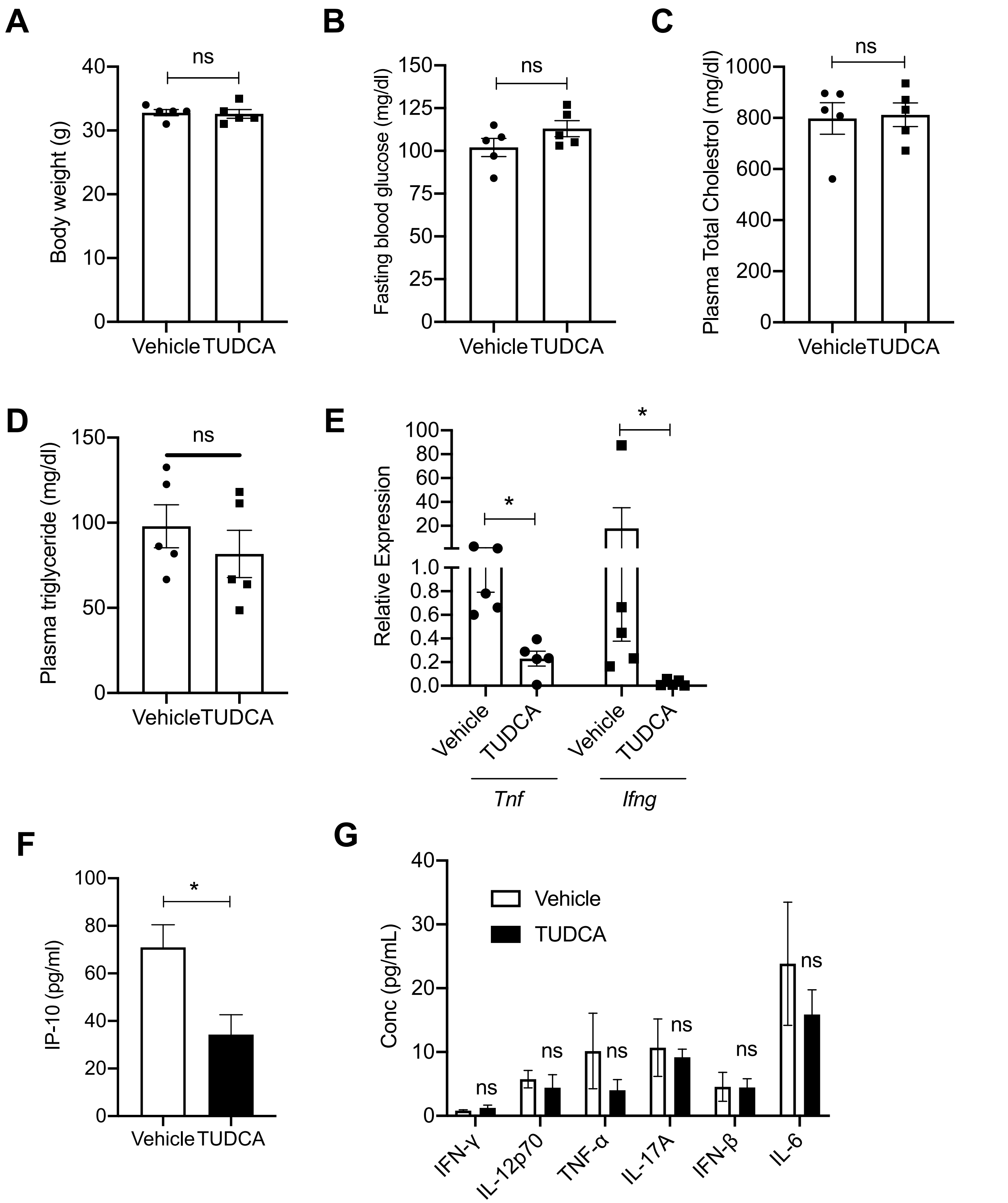


**Supplementary Figure 6.** *Apoe*$^{-/-}$ mice were fed WD for 3 wks alongwith daily intraperitoneal administration of vehicle or TUDCA (150 mg/kg) to appropriate groups of mice as indicated. **(A)** Body weight, **(B)** 5 h-fasting blood glucose, **(C)** plasma total cholesterol, and **(D)** plasma triglyceride was measured and represented as mean  SEM. n = 5 mice per group. ns, no significant difference as measured by Mann-Whitney test. **(E)** RNA isolated from spleen of vehicle or TUDCA-treated mice were analyzed for expression of *Tnf* and *Ifng* by qPCR. **(F and G)** Multiplex-ELISA based analysis of several cytokines in the serum isolated from vehicle or TUDCA-treated mice. n = 5 mice per group. *, $p < 0.05$.
